# Local DNA compaction creates TF-DNA clusters that enable transcription

**DOI:** 10.1101/2024.07.25.605169

**Authors:** Noémie M. Chabot, Ramya Purkanti, Alessia Del Panta Ridolfi, Damian Dalle Nogare, Haruka Oda, Hiroshi Kimura, Florian Jug, Alma Dal Co, Nadine L. Vastenhouw

## Abstract

Transcription factor (TF) clusters have been suggested to facilitate transcription. The mechanisms driving the formation of TF clusters and their impact on transcription, however, remain largely unclear. This is mostly due to the lack of a tractable system. Here, we exploit the transcriptional activation of *mir430* in zebrafish embryos to simultaneously follow the dynamic formation of a large Nanog cluster, the underlying DNA, and transcription output by live imaging at high temporal and spatial resolution. We find that the formation of a Nanog cluster that can support transcription requires local DNA compaction. This brings more Nanog-binding sites into the cluster, and therefore more Nanog. Importantly, we find that Nanog stabilizes this TF-DNA cluster, which emphasizes the interdependent relationship between TFs and DNA dynamics in cluster formation. Once the Nanog-DNA cluster at the *mir430* locus reaches a maximum amount of Nanog, transcription begins. This maximum is a locus-intrinsic feature, which shows that the locus self-regulates the recruitment of an optimal amount of Nanog. Our study supports a model in which endogenous TF clusters positively impact transcription and form through a combination of DNA binding and local DNA compaction.

## Introduction

Transcription factors (TFs) often form clusters in the nucleus ^1–8^. It has been proposed that the local increase in protein concentration in clusters accelerates biochemical reactions ^9^. In the nucleus, for example, TF or co-activator clustering has been demonstrated to decrease TF search time ^10^, increase and stabilize TF binding to DNA ^10^, bring together regulatory elements ^11–13^, and enhance the recruitment of other TFs, co-activators or RNA Polymerase II (RNA Pol II) ^1,6,12–15^. In line with these observations, it has been shown that clustering of TFs can increase the efficiency of transcription ^12,13,15–19^, in some cases by increasing burst frequency ^14,20,21^. Most of these conclusions, however, stem from research on artificially induced clusters ^14–19,21^ or mutated TFs in the context of cancer ^22^. For physiological, non-pathological clusters, data on their effect on transcriptional activity is sparse and conflicting ^12,13,23–26^.

TF assembly into clusters has been proposed to be driven by binding of TFs to DNA, followed by the recruitment of additional factors, potentially facilitated by interactions between intrinsically disordered regions (IDRs) in these proteins ^27,28^. According to this model, the size of clusters would heavily depend on the number of TF binding sites in DNA. Indeed, number and density of TF binding sites has been shown to impact cluster size both *in vitro* and *in vivo* ^21,29^, and the few endogenous sequences that have been shown to seed TF clusters are mostly super enhancers ^5,6,30^, which are characterized by high numbers of TF binding sites ^31^. It thus seems clear that the number of TF binding sites is key in cluster formation. *In vitro* studies, however, have shown that TF clusters can pull in DNA ^32–34^, suggesting a dynamic interaction between TF clustering and the underlying DNA. How DNA and TFs act together to generate TF clusters and regulate transcription *in vivo* however, is not clear.

To understand how clusters form and impact transcription *in vivo*, the dynamics of cluster formation needs to be studied in relationship to the underlying DNA as well as transcriptional output. This has been difficult to achieve in practice, because TF clusters are often small, numerous, and highly dynamic, and in most cases, it is unclear on what sequence they form. In zebrafish embryos, transcription is initially absent after fertilization, and invariably begins with the transcription of the *mir430* locus ^35,36^. This is visible as two large transcription bodies in a nucleus that is otherwise transcriptionally inactive ^2,37–42^. Nanog is essential for *mir430* transcription ^2,43^. It forms multiple clusters in the nucleus, two of which colocalize with *mir430* transcription. Here, we use a live-imaging approach and exploit the transcriptional activation of *mir430* to analyze the formation of a Nanog cluster, how this relates with the organization of mir430 DNA, and how it impacts transcription activation.

## Results

### Nanog clusters associated with mir430 transcription are the largest and brightest

To investigate the relationship between Nanog and *mir430* transcription, we simultaneously visualized Nanog, transcription initiation, and MiR430 transcripts in live embryos. To this end, we injected 1-cell stage zebrafish embryos that lack endogenous Nanog (MZ*nanog*^-/-^) with synthetic mRNA encoding Nanog-mNeonGreen (mNG), Cy5-labelled antigen-binding fragments (Fab) targeting the initiating form of RNA Pol II (RNA Pol II phosphorylated on Serine 5 (RNA Pol II Ser5P)) ^44–47^, as well as an array of fluorescently tagged (Lissamine) antisense oligonucleotides designed to detect MiR430 transcripts (<u>Mo</u>rpholinos for the <u>VI</u>sualization of <u>E</u>xpression, or MoVIE ^38^). We imaged developing embryos on a spinning disk confocal microscope and performed our analysis in 1k-cell embryos, unless otherwise indicated. We observed, as before ^2,41^, that Nanog forms multiple clusters in the nucleus (Fig. 1a, Extended Data Fig. 1). Two of these colocalize with RNA Pol II Ser5P and MiR430 transcript signals. The Nanog clusters that colocalize with MiR-430 transcripts appear to be larger and more intense than other clusters in the nucleus (Fig. 1a). We quantified the volume and intensity of Nanog clusters that do or do not colocalize with the first RNA Pol II Ser5P transcription bodies that appear during the cell cycle (and MiR430 transcripts) and found that the Nanog clusters associated with MiR430 transcripts are indeed the largest and the brightest in the nucleus at the time of *mir430* activation (Fig. 1b, Extended Data Fig. 2).

**Figure 1.**
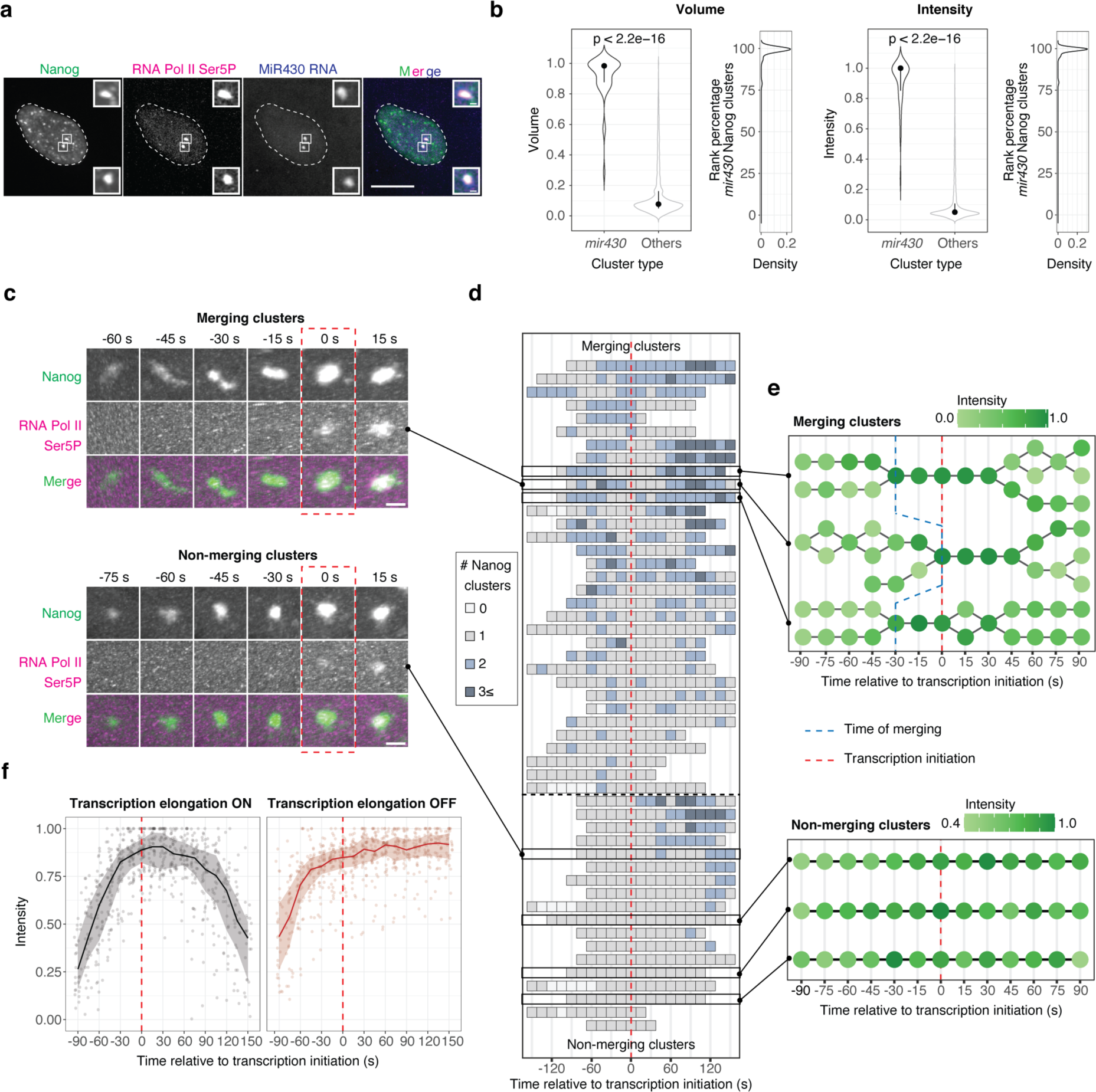
Merging of clusters results in high amount of Nanog associated with *mir430* transcription. **a.** Visualization of Nanog (mNeonGreen; green), initiating RNA Polymerase II (RNA Pol II Ser5P Fab (Cy5); magenta), and *mir430* transcription (MoVIE lissamine; blue) at the 1k-cell stage. Insets are zooms of the two Nanog clusters colocalizing with RNA Pol II Ser5P and MiR430 transcripts. Shown are representative images at the time of *mir430* transcription initiation. Scale bars are 10 and 1 μm (insets). All images represent maximum intensity projections in the z direction. **b.** Violin plots and density plots representing volume (left) and total intensity (right) of Nanog clusters that colocalize with *mir430* transcription (*mir430,* N=3, n=41) or not (others, N=3, n=3781). Density plots represent the rank percentage of the Nanog clusters colocalizing with *mir430* transcription in comparison to other Nanog clusters. Values are normalized to the lowest and the highest values in each nucleus. Pairwise non-parametric Wilcoxon-Mann-Whitney test was performed. **c.** Visualization of Nanog clusters (mNeonGreen, green), and RNA Pol II Ser5P (RNA Pol II Ser5P Fab (Cy5), magenta). Shown are representative images of a merging event and a non-merging event (complete sequence in Supplementary Video 1). Time is relative to transcription initiation. Scale bars represent 1 μm. Images are snapshots from the 3D rendering of the Imaris software. **d.** Schematic representation of all individual Nanog tracks in the dataset (separated by merging and non-merging), centered on transcription initiation (N=5, n=33 for merging clusters; N=5, n=18 for non-merging clusters). The number of Nanog clusters at each time point is indicated by different shades of blue. Tracks are aligned at transcription initiation. Tracks in black boxes are further represented in panels c and e. **e.** Three representative tracks of merging and non-merging Nanog clusters colocalizing with *mir430* transcription, centered on transcription initiation. Shades of green indicate total intensity for each Nanog cluster. Values are normalized to the maximum value for each track. The blue dashed line indicates merging time. **f.** Total intensity of Nanog clusters associated with *mir430* transcription as a function of time (relative to transcription initiation) with (left panel, N=5, n=51) and without (right panel, N=3, n=48) transcription elongation. If the Nanog cluster associated with transcription was the result of a merging event, we summed up the total intensity of all clusters per time point. The bold line represents the median and ribbon the 25^th^ and 75^th^ percentile of the distribution. Values are normalized to the maximum value for each track. In panels c-f, the red dash line/rectangle indicates the transcription initiation time. In this Figure, N is the number of biological replicates, and n the number of clusters.

### mir430 transcription starts when Nanog reaches a maximum

We proceeded to analyze how the Nanog signal associated with MiR430 transcription evolves prior to transcription. We tracked Nanog signal in single nuclei and associated it with the initiation of *mir430* transcription (Fig. 1c, d). We observed that Nanog signal is initially rather diffuse, and that over time, clusters appear. These clusters are highly dynamic and often split and merge as has been observed before for TF clusters (Sabari et al., 2018; Cho et al., 2018; Sharma et al., 2021; Kim et al., 2023; Gaskill et al., 2023). We noticed that in most cases (65%), two or more Nanog clusters merge to form one Nanog cluster prior to transcription (defined as merging clusters; Fig. 1c, d, Supplementary Video 1). In other cases (35%) we see only one cluster prior to transcription start (defined as non-merging clusters; Fig. 1c, d, Supplementary Video 1). For both merging and non-merging clusters, the amount of Nanog in individual clusters increases prior to *mir430* transcription (Fig. 1e). To investigate how the increasing amount of Nanog relates to the onset of *mir430* transcription, we used RNA Pol II Ser5P signal to identify the start of *mir430* transcription and identified the associated Nanog cluster. We then tracked this Nanog cluster back in time to determine how its fluorescence intensity evolved prior to transcription initiation. If the cluster that activates the *mir430* locus was the result of a merging event, we report the combined intensity of these individual clusters. This revealed that Nanog intensity increases steadily prior to transcription, peaks at transcription initiation and decreases afterwards (Fig. 1f, left panel). To investigate the effect of transcription elongation on Nanog intensity, we repeated the experiment in the presence of α-amanitin, which inhibits transcription elongation ^48^. In this case, Nanog intensity does not decrease after reaching a maximum (Fig. 1f, right panel). This is in line with the observation that *mir430*-associated Nanog clusters decrease in intensity after transcription initiation ^41^. Remarkably, however, we observe that even in the absence of transcription, a maximum in Nanog intensity is reached. This suggests that there is an upper limit to the amount of Nanog that can associate with the *mir430* locus. This maximum appears to be a locus-intrinsic feature and not the consequence of limiting amounts of Nanog, because the mean intensity of free Nanog is stable around the time of transcription initiation (Extended Data Fig. 3a). We conclude that the total amount of Nanog at the *mir430* locus increases prior to transcription and that transcription initiates when it reaches a maximum.

### High amount of Nanog can be reached without observable cluster-merging

We next characterized the effect of cluster merging on the amount of Nanog signal by plotting the intensity of clusters that merge prior to *mir430* transcription (Fig. 2a). Here, we report the intensities of individual clusters before merging, and the intensity of merged clusters after merging. As expected, merging increases the total amount of Nanog. Importantly, merging has a negligible effect on the concentration of Nanog in the cluster, but rather increases the volume, and as such the total amount of Nanog in the cluster (Fig. 2a). In agreement with the need to reach a high amount of Nanog for transcription initiation, transcription follows merging in 94% of the cases in which merging can be observed (Fig. 2b). We conclude that high amounts of Nanog at the *mir430* locus can result from the merging of multiple clusters before transcription initiation. Merging of Nanog clusters is, however, not a prerequisite for transcription (Fig. 1c, d). In fact, merging and non-merging clusters show a similar increase in total Nanog intensity prior to transcription as merging ones (Extended Data Fig. 3b), and they initiate transcription at the same time during the cell cycle (Fig. 2c). This shows that the amount of Nanog in non-merging clusters evolves exactly as in merging ones and raises the possibility that merging and non-merging clusters are two representations of the same process. If this were the case, observing just one cluster could mean that merging happened prior to the image acquisition, or the imaging did not reach the temporal resolution to be able to observe it. To investigate this, we analyzed how quickly merging happens by resolving the cases in which we observe merging by the time for which we detect separate clusters (Fig. 2d). This revealed that in most cases (42%), we can see separate clusters for only 15 seconds and only in 18% of the cases we see multiple clusters for more than a minute (Fig. 2d). Thus, merging typically happens very quickly. If merging and non-merging are indeed the same process, it would be predicted that in cases where we observe merging, this is required to reach a high total amount of Nanog. To test this, we compared the increase in the total amount of Nanog between merging and non-merging cases (Fig. 2e). For merging cases, we plotted the amount of Nanog in individual clusters (individual), as well as the sum of individual clusters that merge (sum). Comparing these with the plot for non-merging cases showed that in merging cases, merging is required to reach sufficiently high amounts of Nanog to activate transcription. We conclude that Nanog clusters for which we observe merging and Nanog clusters for which we do not observe merging are different representations of the same process, and that in both cases, the required amount of Nanog to activate *mir430* transcription is reached.

**Figure 2.**
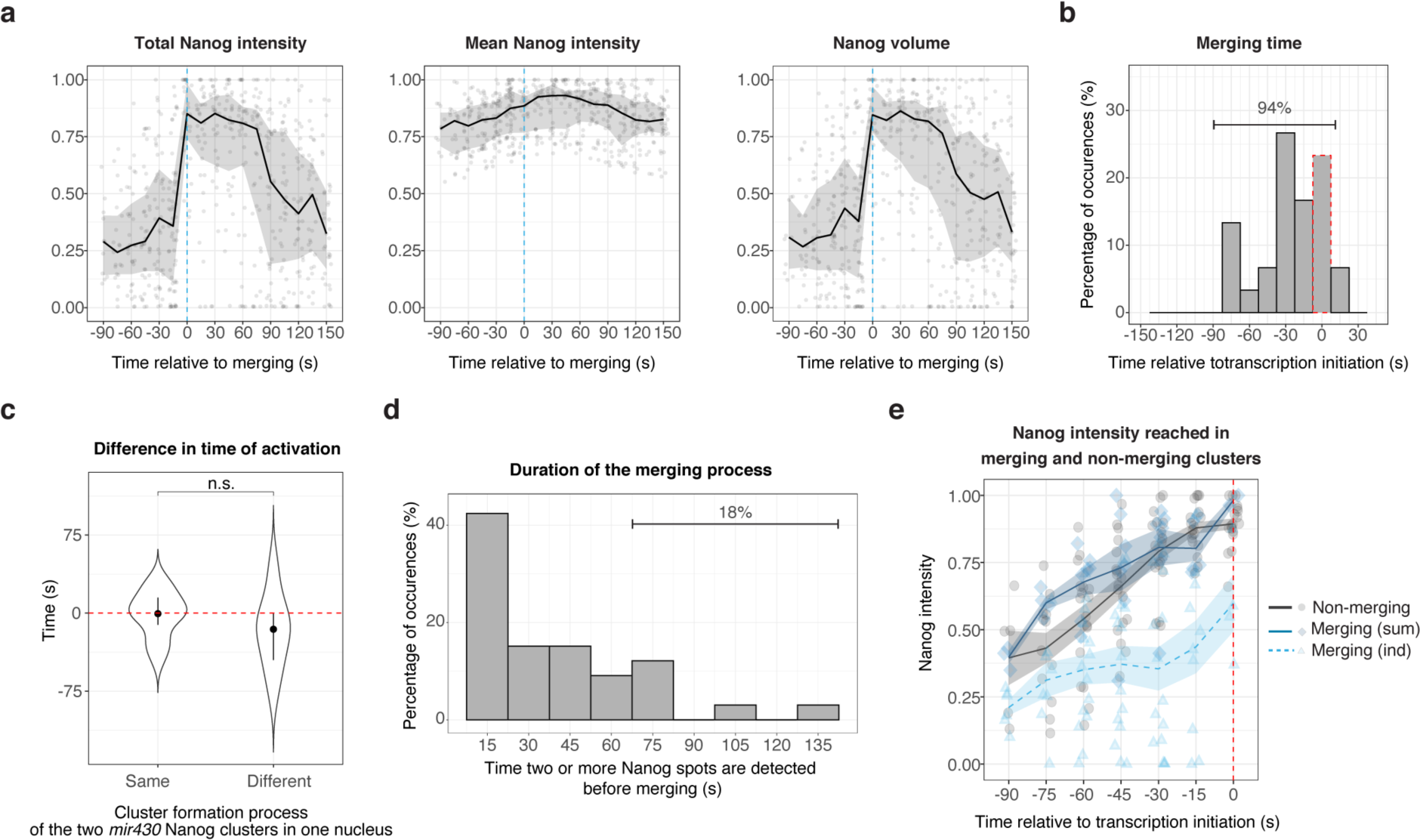
Merging and non-merging cases are functionally identical. **a.** Total fluorescence intensity (amount), mean intensity (concentration), and volume of Nanog signal in merging Nanog clusters (N=5, n=33) associated with *mir430* transcription as a function of time (relative to merging). Here, we report the intensities of individual Nanog clusters before merging, and the intensity of merged clusters after merging. The bold line represents the median and ribbon the 25^th^ and 75^th^ percentile of the distribution. **b.** Histogram showing the time at which Nanog clusters (N=5, n=33) merge relative to transcription initiation. **c**. Difference in time of activation between the two *mir430*-associated Nanog clusters per nucleus, depending on if they are forming the same way (both non-merging or both merging, N=5, n=10) or if they are form differently (one merging and one non-merging, N=4, n=13). Black spots represent the median and vertical bars the 25^th^ and 75^th^ percentile of the distribution. Pairwise non-parametric Wilcoxon-Mann-Whitney test was performed. n.s, indicates p > 0.05. **d.** Percentage of merging clusters for which two or more spots can be observed for the indicated time (N=5, n=33). **e.** Normalized intensity of merging and non-merging Nanog clusters associated with *mir430* transcription relative to transcription initiation. The lines represent the mean of the distribution with the ribbon indicating the standard error of mean. The value of the non-merging clusters is shown as a black solid line (N=4, n=18). The value of merging ones is shown as the sum of the merging clusters (blue solid line, N=5, n=13), as well as individually (blue dashed line, N=5, n=26). For individual clusters, only tracks for which we can detect individual clusters for at least three consecutive time points before merging were plotted. For panels a, b, and e, the red dash line/rectangle indicates the transcription initiation time. In this Figure, N is the number of biological replicates, and n is the number of clusters.

### In vivo labelling of the mir430 locus using a dCas9 approach

To understand how the Nanog clusters that activate *mir430* transcription are spatially related to the *mir430* locus, we adapted a dCas9 labelling approach ^49^ to visualize the DNA of the endogenous *mir430* locus live. We took advantage of the repetitive nature of the locus (Extended Data Fig. 4) and used two single guide RNAs (sgRNAs) that together bind to the locus twenty times^37^ (Extended Data Fig. 4). We inserted eight MS2 loops in the tetraloop of each guide RNA ^50^, and visualized them with MCP protein tagged with mNG. We injected embryos at the 1-cell stage with pre-assembled sgRNA-dCas9 complexes, together with mRNA encoding MCP-mNG (Fig. 3a). This resulted in a clear signal corresponding to the *mir430* locus, as evidenced by the colocalization with MiR430 RNA (Fig. 3b). Injections without dCas9, guide RNAs, or the target locus (*mir430*^-/-^ ^2,42^) resulted in the loss of signal (Fig. 3b), further confirming that our technique detects the *mir430* locus specifically.

**Figure 3.**
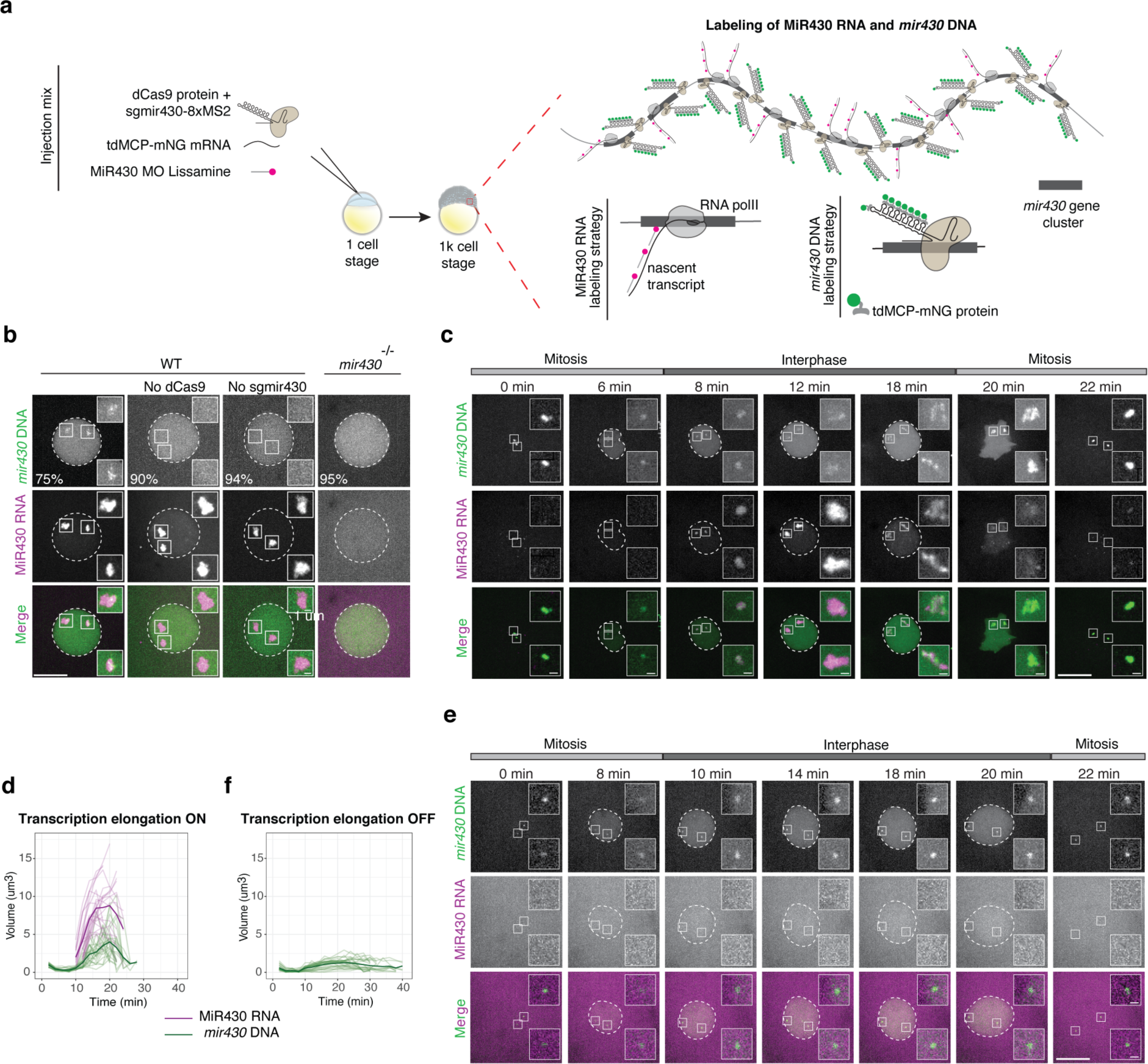
Strategy to specifically visualize the *mir430* locus in living embryos. **a.** Schematic of *mir430* DNA and RNA *in vivo* labelling technique (see Methods). **b.** Visualization of *mir430* DNA (tdMCP-mNG; green) and MiR430 RNA (MoVIE-lissamine; magenta) in WT embryos (+/- dCas9 and +/- sgRNAs), as well as in *mir430* -/- embryos. Shown are representative images of individual nuclei at the midpoint of the cell cycle (from metaphase to metaphase). Scale bars are 10 and 1 μm (insets). Percentages indicate the fraction of nuclei in which we observe the shown phenotype. **c**. Visualization of *mir430* DNA (tdMCP-mNG; green) and MiR430 RNA (MoVIE-lissamine; magenta) in WT embryos. Shown are representative images of individual nuclei during the cell cycle (metaphase to metaphase). Scale bars are 10 and 1 μm (insets). **d.** Quantification of the changes in volume of *mir430* DNA (green) and MiR430 RNA (magenta) signal (N=3, n=24) during the cell cycle (from metaphase to metaphase). Each line is an individual allele, bold lines represent the mean. **e,f**. As c,d but in absence of transcription elongation. For f, N=3, n=28. In b, c and e, images represent maximum intensity projections in the z direction. In this Figure, N is the number of biological replicates, and n is the number of *mir430* alleles.

It has previously been shown that DNA of long, highly expressed genes expands when it is transcribed ^37,39,41,51^. Hence, to test our method, we asked if we could detect an expansion of the long and highly expressed *mir430* locus as it starts to be transcribed. To this end, we followed the *mir430* locus during a complete cell cycle. This revealed a coordination between transcriptional activity and an increase in the volume of the *mir430* locus (Fig. 3c, d). Such expansion of the *mir430* locus was not observed when transcription of the locus was inhibited with α-amanitin (Fig. 3e, f). Thus, our method faithfully detects the *mir430* locus live, and can be used to investigate how the Nanog clusters that activate the *mir430* locus relate to this locus in nuclear space.

### The Nanog clusters that activate mir430 transcription are seeded by the locus itself

We hypothesized, in light of the size of the *mir430* locus ^39,41^ and its high number of Nanog binding sites (Extended Data Fig. 4), that Nanog clusters that ultimately activate *mir430* transcription could form on the locus itself (Fig. 4a). Alternatively, however, they could form away from the locus and subsequently move towards it (Fig. 4a). To distinguish between these two possibilities, we combined the visualization of Nanog (Nanog-HaloTag (JFX650)), the *mir430* locus (MCP-mNG), and MiR430 transcripts (*mir430* MoVIE lissamine) (Fig.4b). We observed that the *mir430* DNA signal is initially rather weak and becomes better visible as we approach transcription initiation. Often, the *mir430* DNA signal is punctuated, which we propose reflects differences in DNA density. As before (Fig. 1c), we observe that Nanog clusters are highly dynamic, and that there are merging and non-merging Nanog clusters. Overall, we observe a high degree of overlap between *mir430* DNA signal and Nanog signal, even before transcription initiation (Fig. 4b). A quantification of how often Nanog clusters overlap with the signal of *mir430* locus, confirmed that most clusters (∼90%) colocalize with *mir430* at the time of merging as well as at all the earlier time points (Fig. 4c, left panel). Because we cannot do this analysis for non-merging clusters, we also used the time of transcription initiation (Fig. 4c, middle and right panels). Both for merging and non-merging clusters, this confirms that most clusters (98% for merging, 97% for non-merging) colocalize with *mir430* at the time of transcription initiation as well as at earlier time points. We conclude that the Nanog clusters that activate the mir430 locus are seeded by the locus itself. Importantly, if merging clusters form on the *mir430* locus, merging does not increase the total amount of Nanog associated with the *mir430* locus, but rather brings all Nanog into one place. This has important implications for our interpretation of the impact of cluster merging on transcription (see discussion).

**Figure 4.**
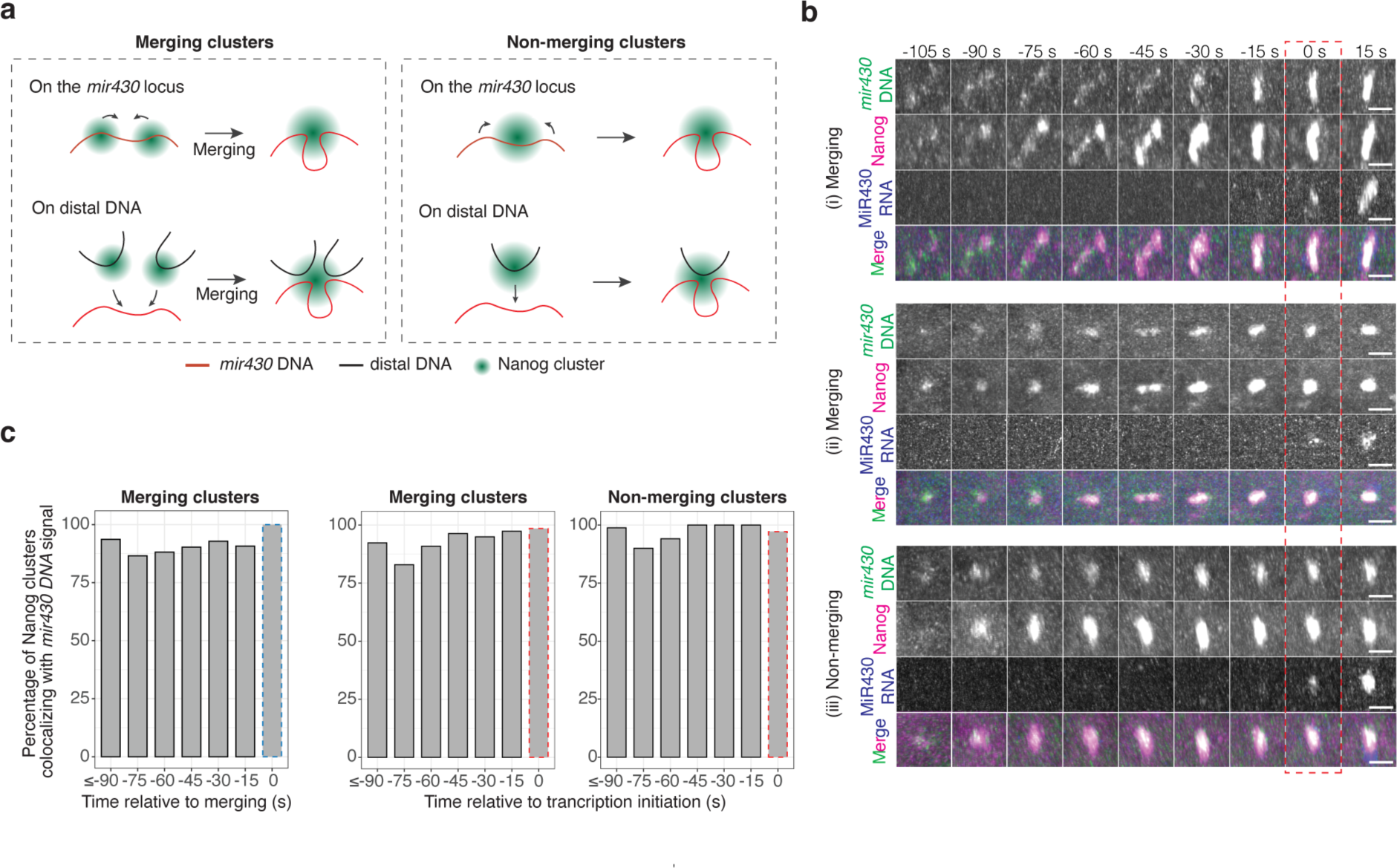
Nanog clusters that activate *mir430* transcription are seeded by *mir430* locus. **a.** Schematics representing two hypotheses concerning the localization of *mir430*-associated Nanog clusters relative to the *mir430* locus. **b.** Visualization of *mir430* DNA (tdMCP-mNG; green), MiR430 RNA (MoVIE-lissamine; blue), and Nanog (HaloTag (JFX650); magenta) in *nanog* -/- embryos. Shown are representative images of timelapse movies showing (i) Merging Nanog clusters (and associated *mir430* DNA) that are separated for a couple of timeframes, then merge (t=-30s) followed by transcription initiation (t=0s); (ii) Merging Nanog clusters (and associated *mir430* DNA) that split (t=-45s) and then merge rapidly (t=-30s), followed by transcription initiation; (iii) Non-merging: a single Nanog cluster (and associated *mir430* DNA) that grows in intensity, followed by transcription initiation. Scale bars are 1 μm. Movies are aligned at transcription initiation, which is boxed in red. All images are snapshots from the 3D rendering of the Imaris software. **c**. The percentage of merging Nanog clusters that colocalize with the *mir430* locus signal for at least one voxel prior to / during merging (left plot; merging; N=4, n=57), and the percentage of merging and non-merging Nanog clusters that colocalize with the *mir430* locus signal for at least one voxel prior to/during transcription initiation (merging N=4 and n=57; non-merging, N=4 and n=32). See Methods for details. The red and blue dashed rectangle indicates transcription initiation. In this Figure, N is the number of biological replicates, and n is the number of tracks.

### Local DNA compaction brings Nanog clusters together

If the Nanog clusters that activate the *mir430* locus are seeded by the locus itself, local DNA compaction could be a potential mechanism to bring Nanog clusters together. To explore this possibility, we analyzed the shape changes of the *mir430* DNA signal in relation to the onset of transcription (Fig. 5a). We segmented the *mir430* locus in 3D and determined the radial distances between the center of gravity and each edge pixel of the maximum-intensity projected mask in 2D for each locus (see Methods). We then calculated the coefficient of variation (CoV) of the radial distances to describe the shape of the locus. According to this metrics, if the locus is fully compacted, the value would be 0.08 (see Methods) whereas higher values would correspond to more elongated shapes (Fig. 5a, left). As such, the CoV of the radial distances within the segmented *mir430* DNA signal can be used as a proxy for its compaction state. Using this approach, we detected a decrease in the CoV before transcription initiation and the lowest value is reached at the time of transcription initiation (Fig. 5a), regardless of the detectability of merging events (Extended Data Fig. 5a). Importantly, the CoV of the radial distances decreases to values measured during mitosis (dashed green line in Fig. 5a), suggesting that the locus is highly compacted. We conclude that transcription starts at the most compacted state of the *mir430* locus.

**Figure 5.**
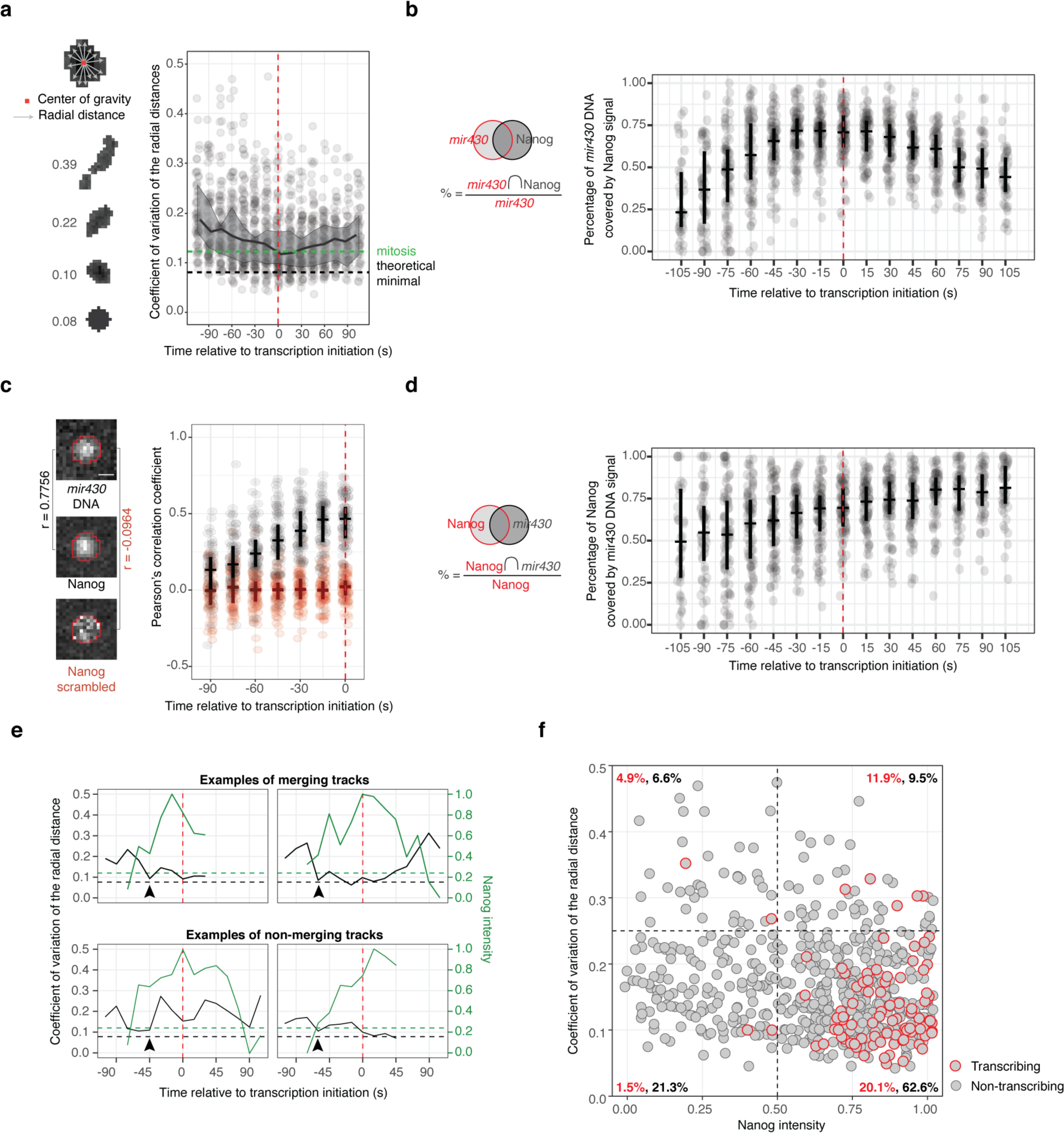
Compaction of the *mir430* locus drives the formation of a Nanog-DNA cluster. **a**. Left: Schematic representation of the approach used to calculate the CoV of the radial distances, as well as examples. Right: The CoV of the radial distances for all tracks is plotted as a function of time, relative to transcription initiation. The black line indicates the median of the distribution and the associated ribbon the 25th and 75th percentile of the distribution. The green dashed line shows the average value of the coefficient of variation of the radial distance of the *mir430* DNA mask measured during mitosis (see Methods). The black dashed line shows the theoretical minimal value for a mask considered as a perfect sphere (see Methods). **b.** Percentage of *mir430* DNA signal that is covered by Nanog signal as a function of time relative to transcription initiation. The vertical lines represent respectively the 25 and 75% of the distribution, while the horizontal line represents the median. **c**. Left: Schematic representation of the approach used to calculate the correlation between *mir430* DNA and Nanog signal. Right: Boxplots showing the Pearson’s correlation score between *mir430* DNA mask and the associated Nanog signal (black) or scrambled Nanog signal (red) for all tracks, relative to transcription initiation. **d.** Percentage of *mir430*-associated Nanog signal that is covered by *mir430* DNA signal as a function of time. The vertical lines represent respectively the 25 and 75% of the distribution, while the horizontal line represents the median. **e.** Four selected tracks (two merging and two non-merging) showing the coefficient of variation of the radial distances of *mir430* DNA mask (black) and associated Nanog total intensity (green), relative to transcription (complete dataset in Extended Data Fig. 6). The green and black dashed lines are as in panel a. Black arrowheads indicate time points for which *mir430* DNA compaction reaches a minimum for the first time. **f.** Scatterplot showing the coefficient of variation of the radial distances of the *mir430* locus (for data points between -105 s until transcription initiation) as a function of the normalized total intensity of Nanog clusters. A red outline indicates associated transcriptional activity. The percentage of *mir430* alleles in each quadrant is indicated in black. The percentage of active alleles in a quadrant as a fraction of all alleles in that quadrant is indicated in red. For all panels, the red dash line indicates transcription initiation. For panels a, b, c, d and f, N=3 biological replicates and n=89 is the number of tracks.

To investigate how compaction of the *mir430* locus relates to Nanog accumulation, we next included Nanog signal in our analysis. First, we observed that prior to transcription activation, *mir430* DNA signal is increasingly covered by Nanog signal (Fig. 5b). We then asked whether the intensity of Nanog signal correlates with the intensity of the *mir430* DNA signal. To do so, we determined the correlation between Nanog and *mir430* DNA intensities within the *mir430* DNA mask. We observe that the correlation increases prior to transcription (Fig. 5c, in black). As a control, we scrambled the pixel intensities of Nanog within the mask. In this case, no increase in the correlation between signals was observed (Fig. 5c, in red). These results show that prior to transcription, the increasing amount of Nanog correlates with the local compaction of *mir430* DNA. In agreement with this observation, we find that the Nanog signal is increasingly covered by the *mir430* DNA signal (Fig. 5d). We note that similar results were obtained independent of whether merging Nanog clusters were observed (Extended Data Fig. 5). We conclude that local DNA compaction brings Nanog and DNA together in a TF-DNA cluster.

Despite the correlation of transcription initiation with *mir430* locus compaction in averaged data (Fig. 5a), plots of individual alleles show that the condensation process is highly dynamic, and transcription does not always begin the first time that the DNA is in its most compacted state (arrowheads in Fig. 5e, Extended Data Fig. 6). To better understand the relationship between *mir430* locus compaction and Nanog accumulation, we added the total amount of Nanog associated with the *mir430* locus to the compaction plots. Taking both parameters into account, we observed that transcription often initiates when DNA is in the most compacted state but only once higher amounts of Nanog have accumulated on the *mir430* locus (Fig. 5e). This supports a model in which local DNA compaction is important but only when there is enough Nanog to be brought together. To look at this in more detail, we plotted the Nanog amount on the *mir430* locus as a function of the compaction state of the *mir430* locus (Fig. 5f). We observe that high levels of Nanog often correlate with compaction, as 62.6% of all observations are found in the lower right quadrant. Of these, 20.1% are associated with *mir430* transcription, which is a higher percentage than in the other quadrants. We conclude that a local compaction of the *mir430* locus helps to create a TF-DNA cluster that facilitates transcription initiation.

### Nanog stabilizes TF-DNA clusters

Our data shows that the Nanog clusters that activate *mir430* transcription are seeded by the *mir430* locus (Fig. 4), and that compaction of the locus facilitates the formation of a Nanog- DNA cluster that enables transcription initiation (Fig. 5). Because it has been shown that TFs can pull in DNA ^32–34^, we next set out to test whether Nanog itself contributes to the local compaction of the *mir430* locus. Here, we used a parameter for DNA compaction that is based on the changes in relative distances between local maxima of intensity in the *mir430* DNA signal (Fig. 6a). We define the locus as compacted when we detect only one density (distance equal to 0), and as decompacted otherwise. To study the compaction of the *mir430* locus in the presence or absence of Nanog independently of transcriptional output, we compared Nanog mutant embryos, in which *mir430* transcription is absent ^2^, to Nanog mutant embryos injected with Nanog and the transcription inhibitor α-amanitin (Fig. 6b, Extended Data Fig. 7a). We found that loci compact and decompact often (Fig. 6b), which is in line with the dynamic behavior of the *mir430* locus that we observed previously (Fig. 4b). We then calculated the speed at which the locus compacts, as defined by the time it takes for the distance between two or more densities to be reduced to zero (Fig. 6a). This speed is not significantly different with and without Nanog (Fig. 6c, Extended Data Fig. 7b), suggesting that Nanog does not directly impact the speed at which the *mir430* locus compacts. In absence of Nanog, however, loci are more dynamic than in its presence (Fig. 6b). This can be seen in number of fluctuations per locus (1.4 on average with Nanog, versus 1.9 on average without, p=0.026). This prompted us to compare the stability of the locus in the presence and the absence of Nanog. Here, we assess stability based on how long the locus spends in a compacted state (Fig. 6a). This revealed that in nuclei with Nanog present, loci spend significantly more time in the compacted state, compared to nuclei in which Nanog is absent (Fig. 6d, Extended Data Fig. 7c). This suggests that Nanog plays a role in stabilizing TF-DNA clusters. If Nanog indeed plays a role in keeping clusters compacted, one would predict that the average distance between DNA signal densities is shorter in the presence of Nanog than in its absence. This is indeed the case (Fig. 6e, Extended Data Fig. 7d). We thus conclude that Nanog does not drive the compaction of the *mir430* locus but stabilizes Nanog-DNA clusters.

**Figure 6.**
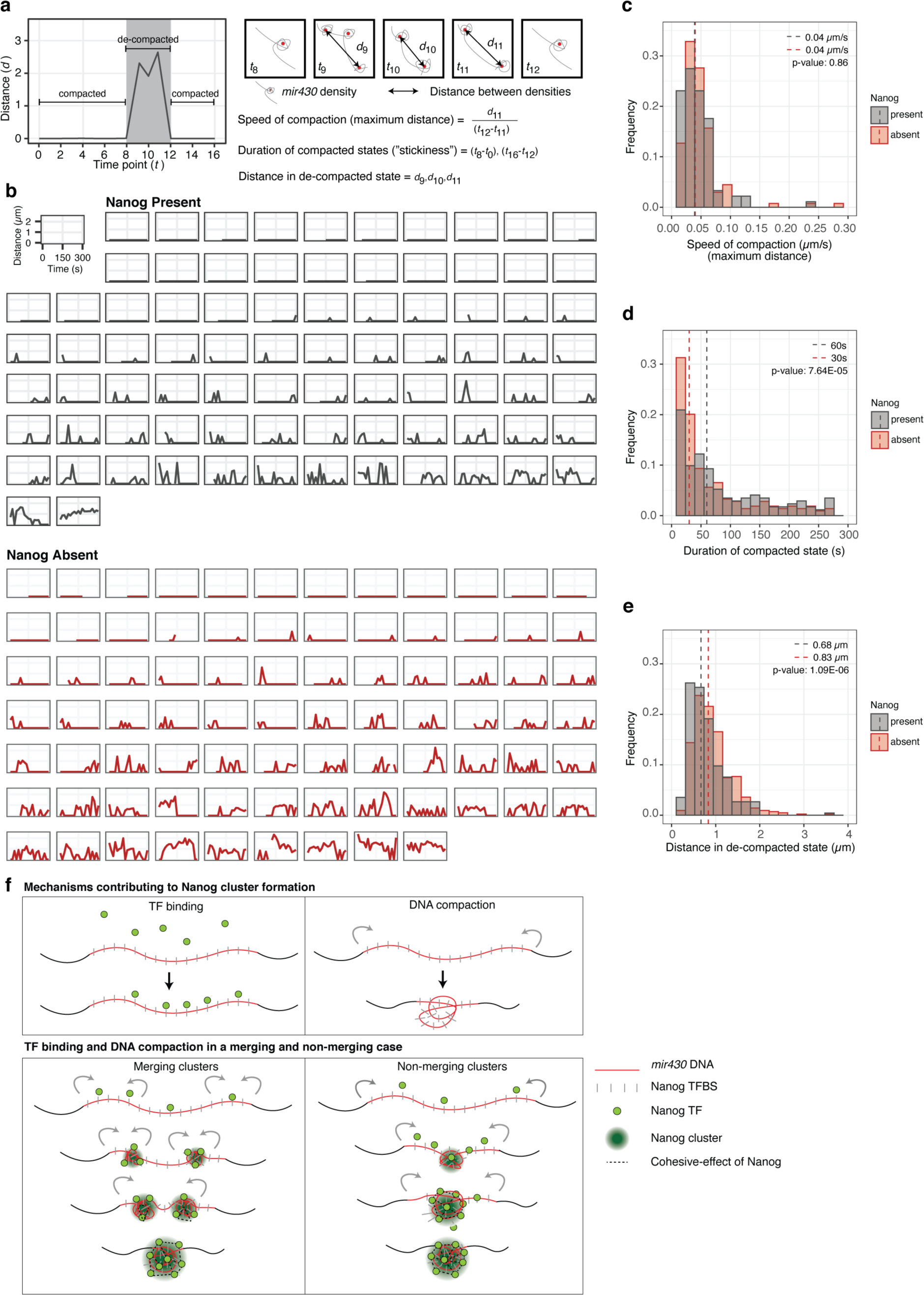
Nanog stabilizes clusters of Nanog and the *mir430* locus. **a. Left: Schematic** representation of the distances between DNA densities, used to derive the parameters assessed in this figure. Right: The gray line represents the *mir430* locus, the local accumulations are detected densities in *mir430* DNA signal. The red dots represent the centroid of detected densities, and the black arrow is the distance between the two most distant densities on the same *mir430* allele (centroid to centroid). **b**. Graphs of the distances between detected densities on the *mir430* DNA channel as a function of time for individual *mir430* DNA alleles in the presence (black) or absence (red) of Nanog. **c-e.** Histograms showing the speed of compaction (from the maximum distance) (c), the duration of compacted states (or stickiness) (d), and the distances in the decompacted states (e) in the presence (black, N=3 and n=80) and absence (red, N=4 and n=81) of Nanog for all stages combined (256-cell stage until High stage). Dashed lines indicate the medians of the distributions. P-values are calculated with one-sided Mann-Whitney test. Values for 1k stage-only are depicted in Extended Data Fig. 7. **f.** Model: Nanog accumulation and DNA compaction together result in the formation of a TF-DNA cluster that enables transcription, in merging as well as non-merging cases. In brief, while Nanog binds to the *mir430* locus, which is rich in Nanog binding sites, locus compaction brings in more Nanog binding sites, and thus more Nanog. Nanog helps to stabilize the resulting TF-DNA cluster. In this Figure, N is the number of biological replicates, and n is the number of *mir430* alleles.

## Discussion

Here, we used a live-imaging approach to study the formation of endogenous Nanog clusters on the *mir430* locus and relate this to transcriptional activity. We find that *mir430* transcription begins when a maximum amount of Nanog is reached. This maximum is reached by binding of Nanog to the locus, in combination with local DNA compaction, which brings the locus and Nanog together in a TF-DNA cluster. Nanog itself does not drive the process of DNA compaction, but rather stabilizes Nanog-DNA clusters.

### Endogenous clusters self-regulate to reach optimal amount of transcription factor

We show that *mir430* transcription begins when a maximum amount of Nanog is reached in the cluster associated with the *mir430* locus. This is in line with previous work in which clustering of transcriptional machinery was shown to enhance the transcription of target genes ^12–15,17–19,21,23,25,26^. Other studies have, however, reported a negative effect of clustering on transcription ^24,52–54^. When negative effects were observed, cluster formation was often manipulated by changing features of the clustered proteins ^52–54^. Because negative effects have been ascribed to molecular crowding ^24,53^ and the precise composition of clusters impacts transcription ^24,52,53^, artificial induction of clustering or changing protein-features can probably explain these observed negative effects of clustering. In our work, we focused on a cluster that forms on an endogenous locus and found that the amount of Nanog that triggers *mir430* transcription, is – at the same time –the maximum amount of Nanog that can be associated with the locus. This suggests that the *mir430* locus regulates the amount of Nanog that can be associated with it. We conclude that there is an optimum amount of TF to activate transcription, and that endogenous clusters can self-regulate to reach this optimum.

### A role for DNA compaction in TF cluster formation

We identified two processes that contribute to the accumulation of sufficient amounts of Nanog to activate *mir430* transcription (Fig. 6f). On the one hand, Nanog binds in high amounts to the *mir430* locus, rich in Nanog binding sites. This has been observed before by us and others ^2,13,41,55^ and is in line with the density of Nanog binding sites on the locus ^39,41^. We find that Nanog binding on the locus often results in the formation of Nanog clusters. On the other hand, the *mir430* locus locally compacts, which brings the *mir430* locus and the associated Nanog together to generate a larger TF-DNA cluster. We note that compaction brings more Nanog into one place, regardless of whether separate clusters visibly merge. Because loading of Nanog and DNA compaction occur simultaneously, it is difficult to identify the precise role of compaction in transcription activation, especially because the total amount of Nanog on the locus does not increase because of compaction. Given the emergent properties of clusters, we can imagine that the formation of a cluster allows for a higher amount of Nanog to be associated with the *mir430* locus than could be achieved simply by Nanog binding to a locus that is not compacted. Along the same lines, a cluster may facilitate the recruitment of additional factors. More detailed biophysical experiments will be needed to determine exactly which emergent properties of TF-DNA clusters impact transcription activation.

### Nanog stabilizes TF-DNA clusters

Our work shows that Nanog stabilizes clusters of locally compacted DNA and Nanog itself. Previous *in vitro* work has shown that TF clusters are able to pull in DNA ^32–34^. We show here, however, that Nanog does not affect the speed by which DNA compacts, but rather the stability of the cluster once formed. Further work will be required to determine which factor(s) drive the compaction itself. TF clustering is facilitated by the presence of IDRs as these mediate low affinity interactions between proteins ^12,56–58^. Specifically, for Nanog, we have previously shown that at least one IDR is required to form clusters at the *mir430* locus ^2^. It is therefore possible that the stabilizing function of Nanog is related to its ability to form clusters, but we cannot exclude the possibility that other factors are involved. Because transcription of the *mir430* locus depends on Nanog ^2,43^ and DNA compaction helps to bring in the required amount of Nanog (this study), we propose that the stabilizing function of Nanog helps to maintain high enough levels of Nanog associated with the *mir430* locus to trigger transcription. Nanog is a well-known pluripotency factor ^59^ with important roles in reprogramming ^60,61^, lineage specification ^62–64^, and zebrafish genome activation ^43,65–67^. A role for Nanog in stabilizing TF-DNA clusters, however, was not reported before, and this is a novel function for Nanog.

## Acknowledgements

We thank members of the Vastenhouw lab (especially Gilles Willemin and Edlyn Wu) for their support, helpful feedback, and stimulating discussions. We thank Shivali Dongre, Maria Cristina Gambetta, Maciej Kerlin, Arianna Penzo, and Aleksandar Vještica for comments on the manuscript, and the following facilities and services for their support: MPI-CBG - fish facility, light microscopy facility, and scientific computing, UNIL - cellular imaging facility, fish facility, EPFL – BioImaging and Optics Core and fish facility. Research in N.L.V’s laboratory was supported by the Max Planck Society, the University of Lausanne, a European Research Council Consolidator Grant (101003023), the Volkswagen Foundation (94773), and the German Research Foundation (VA 1209/2-1). Research in H.K.’s laboratory was supported by the Japan Society for the Promotion of Science KAKENHI (JP18H05527, JP21H04764 and JP20K06484).

## Author contributions

Conceptualization: N.M.C., R.P., N.L.V. Methodology: N.M.C., R.P., H.O. H.K. Investigation: N.M.C., R.P. Data analysis: N.M.C., R.P., A.D.P.R., D.D.N, F.J., A.D.C Writing – original draft: N.M.C. and N.L.V. Writing – reviewing & editing: all authors. Funding acquisition: N.L.V., H.K. F.J. Supervision: N.L.V.

## Data and materials availability

Raw imaging data are available upon request. All other data is available in the main text or the extended data.

## Methods

### Zebrafish handling and molecular biology approaches

#### Zebrafish maintenance and manipulation

Zebrafish were maintained and raised under standard conditions, and according to Swiss regulations (canton Vaud, license number VD-H28). To identify *nanog*^-/-^ fish, we fin-clipped adults and genotyped them as previously described ^68^. Wild type (ABTL), *mir430*^-/-^ or *nanog^-^*

*^/-^* embryos were collected less than 10 minutes after fertilization. We always injected at the 1- cell stage and into the cell. The chorion was either mechanically removed with forceps, or chemically by incubating embryos for 3 minutes in 1.5 mg/mL Pronase E (Sigma-Aldrich, 107433) in Danieau’s 0.3X. To rescue *nanog*^-/-^ embryos, we injected 120 pg of full-length Nanog as before ^66^ or the molar equivalence of this when injecting Nanog fusion constructs. Lissamine-labelled anti-MiR430 morpholino was injected at 25 fmole/embryo ^38^. Nanog was visualized using Nanog-mNG or Nanog-HaloTag, and the mRNAs encoding these were injected at 180 and 210 pg/embryo, respectively. To inhibit transcription, α-amanitin (A2263, Sigma-Aldrich) was injected at 0.25 ng/embryo. Transcription inhibition was confirmed by a developmental arrest prior to gastrulation as described before ^2^. To label RNA Pol II Ser5P, Fabs (αRNA_PolII_Ser5P_Cy5) were injected at 1.8 ng/embryo. To label the *mir430* DNA locus, dCas9 protein was injected at 0.25 ng/embryo, sgRNAs (as a equimolar mix of *sgmir430_1* and *sgmir430_2*, see below) ^37^ were injected at 50 fg/embryo, and tdMCP-mNG mRNA at a 25 pg/embryo. After injection, embryos were raised at 28°C in Danieau’s 0.3X or blue water until the desired stage ^69^. For Nanog-HaloTag labelling, embryos were soaked for 20 minutes in 5 μM of JFX650-HaloTag dye (CS315109, Promega) diluted in Danieau’s 0.3X.

#### Generation of Nanog-HaloTag and NLS/NIS-tdMCP-mNeonGreen

To generate the pCS2-Nanog-HaloTag plasmid, we obtained the HaloTag sequence from the Protein Expression and Purification facility (MPI-CBG) and amplified it by PCR using HaloTag specific primers that added a FseI and an AscI site at the 5’ end. HaloTag was then cloned into an empty pCS2+ vector using Gibson Assembly (E2611, NEB), as well as a sequence encoding a linker protein (5’-GGATCCGCTGGCTCCGCTGCTGGTTCTGGC-3’) ^70^. The Nanog coding sequence was then amplified from a pCS2_Nanog_mNeonGreen plasmid^2^ with primers that added Fse I and Asc I sites and cloned into the pCS2_HaloTag plasmid using T4 DNA ligase (NEB, M0202S). The tdMCP gene was amplified from plasmid pME-NLStdMCP-tagRFP (AddGene #86244) and cloned into pCS2+ plasmid with a zebrafish-codon-optimized mNeonGreen tag at its C-terminus. Sequences for a Nuclear Localization Signal (NLS) and a Nucleus Export Signal (NES) were introduced at the N-terminus.

#### mRNA production

Nanog-mNeonGreen, Nanog-HaloTag and NLS/NIS-MCP-mNeonGreen were *in vitro* transcribed using the mMESSAGE mMACHINE^TM^ SP6 Transcription Kit (AM1340, Invitrogen^TM^) from NotI-linearized plasmids. This was followed by digestion of the DNA template using TURBO DNase (AM1340, Invitrogen^TM^) for 15 min at 37°C. Synthetic transcripts were recovered using the RNeasy MinElute Cleanup Kit (QIAGEN, 7404), and quantified using the NanoDrop (NanoPhotometerⒸ NP80) and Qubit (Qubit fluorometerⒸ, Invitrogen) systems. Size and integrity were verified by gel electrophoresis. Single-use aliquots were stored at -80°C.

#### Recombinant dCas9 expression and purification

The gene encoding a catalytically inactive Streptococcus pyogenes Cas9 (D10A/H840A) (dCas9) was cloned in a T7 expression vector with a His-maltose binding protein (MBP) tag at the N-terminus. The dCas9 sequence also contained two copies of Nuclear Localization Signal (NLS) sequence at the N-terminus and one at the C-terminus, to facilitate nuclear import. The S. pyogenes Cas9 D10A/H840A mutant was expressed in T7 express strain (NEB, C2566H) containing the pRARE plasmid (Novagen, 71405) and cultured at 37°C in terrific broth medium supplemented with chloramphenicol (17 µg/mL), kanamycin (100 µg/mL) to OD_600_ = 0.5. Cultures were then shifted to 18°C and induced with 0.2 mM IPTG overnight. Cells were lysed in a lysis buffer (50 mM Tris pH 8.0, 1M NaCl, 1 mM DTT), supplemented with protease inhibitor cocktail (Roche). To remove any nucleic acid contaminants, polyethylenimine (PEI) was added to the clarified lysate (0.25% w/v) and the sample was clarified by high-speed centrifugation after 10 min incubation on ice. Clarified lysate was filtered through an 0.45-µm filter and loaded on a MBP Trap column. The column was washed with a lysis buffer without DTT and cleavage buffer (20 mM HEPES, 250mM KCl, 10% glycerol, 1 mM DTT). Protein was eluted with elution buffer (20 mM HEPES, 250mM KCl, 10% glycerol, 1 mM DTT and 10mM Maltose) and cleaved with PreScission protease overnight to remove the His-MBP affinity tag. After cleavage, the protein was separated from MBP using cation-exchange chromatography with a 5 ml SP Sepharose HiTrap column (GE Life Sciences). Fractions containing dCas9 protein were pooled and the protein was concentrated with spin concentrators (Amicon Ultra 15, MWCO 30 k; Millipore), diluted to final concentration of 2.5mg/mL using storage buffer (20mM HEPES, 250mM KCl pH 7.25), flash-frozen in liquid nitrogen and stored at -80°C.

#### Preparation of in vitro transcribed gRNAs

The sgRNAs were made by *in vitro* transcription for which the DNA templates were prepared by PCRs on plasmid pPUR-hU6-sgRNA-Sirius-8XMS2 (Addgene #121942) as the template encoding the optimized tracr RNA sequence with the integrated MS2 stem loops^50^. The forward primers were designed uniquely for each sgRNA with an overhang containing T7 promoter, the seed sequence, and a sequence complementary to the plasmid (sg*mir430-*1_F: 5’-taatacgactcactataGAGGGTACCGATAGAGACAA<u>gtttgagagctactgccatgagga</u>-3’ and sg*mir430-*2_F: 5’-taatacgactcactataGGCTGAGTGTTAACGACTG<u>gtttgagagctactgccatgagga</u>-3’). The reverse primer (5’-AAAAAAAGCACCGACTCGGTGCC-3’) was the same for both reactions.

PCR products were purified using QIAquick PCR purification kit (Qiagen, 28104). The purified product was used as a template for T7 *in vitro* transcription (HiScribe T7 High Yield RNA Synthesis Kit (NEB, E2040S). *In vitro* transcribed sgRNAs were DNAse-treated, purified by phenol:chloroform:isoamyl alcohol (25:24:1) extraction, followed by ethanol precipitation. The sgRNA pellets were dissolved in 20 mM HEPES (pH 7.5) and 300 mM KCl. To refold purified sgRNAs, the sgRNAs were incubated at 70°C for 5 min and slowly cooled down to room temperature. MgCl2 was then added to 1 mM final concentration and the sgRNA samples were incubated at 50°C for 5 min and slowly cooled down to room temperature. The sgRNAs were quantified, aliquoted to single-use aliquots and stored at -80°C.

#### Preparation of the mir430 DNA labelling reagents

To prepare reagents for mir*430* locus live visualization, 1-μL aliquot of dCas9 protein (2.5mg/mL) was diluted in 9 μL of 20 mM HEPES, 300 mM KCl solution to a final concentration of 0.25mg/mL. 50 ng of each guide (sg*mir430_*1-8xMS2 and sg*mir430_*2-8xMS2) were mixed with the dCas9 protein solution and incubated at 37°C for 10 minutes. After incubation, the assembled dCas9-sgRNA RNP complex was stored on ice until injections.

#### MiR430 RNA labelling

*pre-miR430* transcripts were visualized using the <u>Mo</u>rpholino <u>VI</u>sualization of <u>E</u>xpression (MoVIE) method as described before ^38^. Briefly, a morpholino oligonucleotide complementary to the 5’ end of *pre-miR430* transcripts is coupled at the 3’ end with the red-emitting chemical lissamine fluorophore (GeneTools, https://www.gene-tools.com/).

#### Preparation of RNA Pol II Ser5P antigen-binding fragments

RNA Polymerase II was visualized using Fab-based live endogenous modification labelling (Fab) conjugated with a Cy5 dye ^45–47,71^. Fluorescently labelled Fabs specific to RNA Pol II Ser5P were prepared from monoclonal antibodies specific to RNA Pol II Ser5 phosphorylation (CMA605/Pa57B7). Purified mouse IgG was digested with Ficin (ThermoFisher Scientific) or Papain (ThermoFisher Scientific), and Fabs were purified using HiTrap protein A-Sepharose columns (GE Healthcare) to remove Fc and undigested IgG. Fabs were concentrated up to more than 1 mg/mL using 10 k cut-off filters (Amicon Ultra-0.5 10 k; Merck), according to the manufacturer’s instruction.

Fluorescent dye conjugation was conducted using 50 or 100 μg of purified Fab fragments. In a typical reaction, 50 μg of purified Fab was diluted in 45 μL PBS, mixed with 5 μL 1 M NaHCO_3_ (pH 8.3) and then with 0.5 μL of Cy5 N-hydroxysuccinimide ester (10 mg/mL in DMSO; cytiva, PF11A25001). After incubating for 1.5h at room temperature with gentle rotation in the dark, unconjugated fluorescent dye molecules were removed using a PD MiniTrap G-25 column (Cytiva, 28918004; pre-equilibrated with PBS). The reaction mixture (50 μL) was applied onto a column and 550 μL PBS was applied; the flowthrough fraction was discarded. Dye-labelled Fab fragments were eluted with 500 μL PBS and concentrated to ∼1.2 mg/mL using a 10-kDa cutoff Amicon Ultracell Centrifuge Filter Unit (Merck, UFC5010BK). Fab concentration and Dye:Fab ratio was measured using a Nanodrop (NanoPhotometerⒸ NP80). Dye:Fab ratios were between 0.6:1.1 and 1:1. Aliquots of labelled Fabs were bead-loaded into HeLa cells to validate that they distributed as expected and were then stored at 4°C in the dark.

### Identification of Nanog binding sites

The Nanog binding motif was determined using ChIP-seq published data ^55^. From this canonical binding motif, all potential binding motifs were uncovered at the *mir430* DNA locus sequence (GRCz11 genome assembly) with the AME software using a 5% false-negative cut-off.

### Imaging

#### Preparing embryos for live-imaging

Mounting was performed using 0.8% low-melting agarose solution (UltraPure Low Melting Point Agarose, 16520050, ThermoFisher) diluted in Danieau’s 0.3X and containing 25% v/v OptiPrep density gradient medium (D1556, Sigma-Aldrich). The agarose solution was melted at 70°C and then kept at 37°C during embryo mounting. Between 10 and 15 embryos were transferred into a glass vial filled with the agarose solution and then moved to the surface of a µ-Dish 35 mm, high imaging dish (ibidi, 81156). After waiting 10 minutes to allow the agarose to polymerize, the plate containing the embryos was brought to the microscope under light-protected conditions.

#### Live-Imaging on the confocal spinning-disk microscope

Imaging was performed on an inverted Nikon Ti2 microscope associated with a Yokogawa CSU-W1 spinning-disk confocal unit using a Nikon 100X Oil CFI Plan Achromat Microscope Objective. Images were acquired using a Photometrics Prime 95B and fluorophores were excited using one of the four available laser lines: 405, 488, 561 and 638. Embryos were maintained at a temperature of 28°C using a fully enclosed temperature-controlled chamber. Most of the imaging was done using simultaneous acquisition using a duo camera system, with one exception: when Nanog, *mir430* DNA and MiR430 transcripts were imaged, Nanog and *mir430* DNA were acquired first, followed by MiR430 transcripts.

### Image processing and analysis

#### Time-lapse max projection, mapping and cropping of nuclei

Each time-lapse was projected on the Z axis using Fiji ^72^ to obtain 2D max-projected images in the Z plane. All time-lapses (recorded in the .nd2 file format) were converted into .ims files using the Imaris file converter (RRID:SCR_007370, Bitplane). If the time-lapse contained several cell stages, these were isolated. Each individual nucleus was assigned a unique ID. As such, any nucleus used for analysis can be traced back to the raw data. Single nuclei were cropped manually using either Fiji or the 3D crop function in Imaris (RRID:SCR_007370, Bitplane).

#### Developmental stage determination

Developmental stage was determined based on the time of mounting of embryos, the time that the embryos were allowed to develop during imaging, and the distances between nuclei. For the latter, we measured the distances between the center of neighboring nuclei in the same focal plane as the inter-nuclear distance in early stages is highly stable across embryos.

#### Correction of channel registration

Shifts in alignment between the two cameras were most often corrected at the software level using a calibration slide. If data still displayed registration shifts between two channels afterwards, such shifts were corrected post-imaging. To this end, nuclei were segmented in both channels, and X, Y, and geometric centered positions were retrieved. Using Imaris, registration was corrected by aligning the geometric center in X and Y.

#### Segmentation of Nanog clusters, mir430 DNA and MiR430 RNA in 3D

Objects were segmented using the following algorithms and parameters in Imaris:

|  | **Nucleus** | **Nanog** | **MiR430**<br>**RNA**<br>**signal** | **Ser5P-Cy5** | ***mir430***<br>**DNA**<br>**signal** | ***mir430***<br>**DNA**<br>**signal** |
| --- | --- | --- | --- | --- | --- | --- |
| **Algorithm** | Surface | Spot | Surface | Surface | Surface | Spot |
| **Channel** | Nanog/*m*<br>*ir430*<br>DNA | Green/Far<br>red | Red | Far red | Green | Green |
| **Initial**<br>**parameters** | Object-Object<br>Statistics | Object-Object<br>Statistics | Object-Object<br>Statistics | Object-Object<br>Statistics | Object-Object<br>Statistics | Object-Object<br>Statistics |
| **Diameter** | 9 µm | XY: 0.325<br>µm<br>Z: 0.650<br>µm | 0.5 µm | 0.325 µm | 0.5 µm | XY:<br>0.325 µm<br>Z: 0.650<br>µm |
| **Smoothing** | 0.5 µm | / | 0.220 µm | 0.5 µm | 0.5 µm | / |

| Thresholding | Variable | Variable | Variable | Variable | Variable | Variable |
| --- | --- | --- | --- | --- | --- | --- |
| Final filtering | Remove other nuclei or vesicles | Remove extranuclear objects | Remove extranuclear objects | Remove extranuclear objects | Remove extranuclear objects | Remove extranuclear objects |
| Tracking | None | Manual | Manual or autoregression model (see below) | Manual or autoregression model (see below) | Manual or autoregression model (see below) | Manual |

#### Manual tracking

After segmentation using the spot algorithm in Imaris (see above for details), Nanog clusters that colocalize with *mir430* transcription were identified using RNA Pol II Ser5P or MiR430 transcripts signal at transcription initiation. Once identified, the Nanog clusters were tracked manually using the manual tracking option in Imaris. For time points before transcription initiation, tracking was performed going back in time, frame by frame, from transcription initiation. All Nanog clusters that clearly connected across time points were manually associated together to the same track. If an association between clusters at consecutive timepoints was not clear, the Nanog track was removed from the dataset. For time points after transcription initiation, all Nanog clusters that colocalized with the transcription body (labeled by RNA Pol II Ser5P or MiR430 transcript signal), were considered as part of the same Nanog cluster and associated to the same track. For *mir430* DNA (segmented with the spot algorithm), spots were manually linked to the same track using the manual tracking option in Imaris. If the *mir430* DNA signal at one time point was detected as more than one spot, spots were linked to the same track.

#### Semi-automated tracking

After segmentation using the shape algorithm in Imaris (see above for details), MiR430 transcripts, RNA Pol II Ser5P or *mir430* DNA signals were tracked automatically using the tracking plugin of Imaris. The algorithm used was the autoregression model, with a maximum connecting distance of 4 μm. The accepted number of missing time points was between 0 and 5 depending on the time in the cell cycle. In cases where automatic tracking was not able to correctly link the same cluster over time, objects were tracked using the manual tracking option in Imaris.

#### Colocalization and Pearson’s correlation score analysis in 3D

Nanog and *mir430* DNA signal were segmented as described above. Once segmented, a 3D mask of the signal was obtained, and each *mir430* allele was isolated. In general, if the segmented masks for Nanog or *mir430* DNA were composed of two or more parts (for example in the case of Nanog merging clusters), the two masks were merged. To calculate the percentage of cases where Nanog and *mir430* DNA signals overlap, we considered both signals as colocalizing if at least one voxel was shared between the two masks. To calculate the percentage of overlap between Nanog clusters and *mir430* DNA, we used a custom MATLAB script that calculated the percentage of voxels in the Nanog mask that overlapped with voxels from the *mir430* DNA mask over the total number of voxels contained in the *mir430* DNA mask. Conversely, to calculate the percentage of overlap between *mir430* DNA and Nanog, we calculated the percentage of voxels in the *mir430* DNA mask that overlapped with voxels from the Nanog mask over the total number of voxels contained in the Nanog mask. We calculated the Pearson’s correlation score between Nanog and *mir430* DNA using the raw pixel intensities of these two signals within the mir430 DNA mask. As a control, the Nanog raw intensity values were scrambled.

#### Determination of the radial distances inside mir430 DNA mask

DNA signal was segmented using the shape algorithm in Imaris (see above for details) then tracked in 3D using TrackMate v7.11.1 ^73^. Object detection was performed using the mask detector and linking using the Advanced Kalman Tracker. Tracks were manually validated and corrected in napari (napari contributors, 2019 at https://zenodo.org/records/8115575). Following tracking, each individually tracked 3D mask was projected into 2D using maximum projection. A 2D convex hull was created around each mask and an array containing the distance from the center of gravity of the convex hull (calculated with subpixel accuracy) to the center of each boundary pixel was calculated for each time frame. This analysis was performed in python using the numpy (1.23.1), skimage (0.19.3) and pandas (1.5.3) libraries. From this array of radial distances, the coefficient of variation (CoV) was calculated for each DNA mask at each time point. To determine the time of transcription initiation, we used the segmented mask of the RNA channel, generated as described above. We considered the initiation of transcription to be the first appearance of non-zero pixels in this masked image. Once an RNA mask was detected, it was associated with the nearest DNA mask to determine the unique onset of transcription per allele. To obtain the theoretical minimal value of the CoV of the radial distances, we generated a perfect circle as a pixelated image with a radius of 4 pixels using the OpenCV circle function. Radial distances were calculated the same way as for *mir430* DNA masks. To calculate the CoV of radial distances value during mitosis, we analyzed *mir430* DNA masks in nuclei with a sphericity less than 0.6 (calculated by Imaris).

#### Association of Nanog with RNA Pol II Ser5P transcription bodies

To associate Nanog with RNA Pol II Ser5P transcription bodies, we used two approaches. First, we calculated the distance in 3D between the gravity center of segmented Nanog clusters and transcription bodies using the Imaris software (*Shortest distance to Surfaces*). Next, we calculated the distance based on the coordinates of the objects from Imaris using R. Only if the calculated distance between the center of gravity of both segmented signals was less than 0.5 μm in Imaris and less than 1 μm in R, Nanog was associated with the RNA Pol II Ser5P transcription bodies.

#### Association of Nanog clusters with MiR430 transcription bodies

To associate Nanog clusters and MiR430 transcription bodies, the same method was applied as described above for Nanog clusters and RNA Pol II Ser5P transcription bodies, with one difference: because the Nanog and MiR430 transcription body signals were acquired with a lag of approximatively 7 seconds, we allowed larger distances. Here, Nanog was associated with the MiR430 transcription bodies only if the calculated distance between the center of gravity of both segmented signals was less than 1 μm in Imaris and less than 1.5 μm using R. Note that in Figure 1, the Nanog clusters closest to *mir430* transcription were considered as mir430-associated Nanog clusters.

#### Categorization of Nanog clusters into merging/non-merging categories

Categorization of Nanog clusters in the merging or non-merging category was performed using R as follows: If Nanog cluster tracks showed a single cluster at all time-points prior to transcription activation it was called a non-merging cluster. If in the last 10 time points before transcription initiation, at least two spots were detected in at least one time point, it was called a merging cluster.

#### Determination of the time of merging

For merging cases, the time of merging was defined as the time when only one Nanog cluster was detected in a time window of -75 to +45 seconds around transcription initiation. If more than one time point met this criterion, the closest time point to transcription initiation was considered as the time of merging, with priority being given to a timepoint before transcription initiation. If no time point with a single cluster was detected in a range of -75 to +45 seconds around transcription initiation, the Nanog clusters were considered as never merging and no time of merging was calculated.

#### Classification of cell cycle phase (mitosis or interphase)

The phase of the cell cycle was determined by features such as the roundness of the nucleus, chromosome compaction, and the distance between two daughter cells. These features were observed using Nanog-mNG, Nanog-HaloTag or MCP-mNG signals.

#### Analysis of compacted and decompacted state for mir430 DNA

##### Time-series analysis of *mir430* loci distances

To determine distances between DNA densities, Mir430 DNA densities were segmented using the spot algorithm (see above for details). If only one spot was detected, we considered the distance to be zero. For all other time points, we calculated the distance between all detected spots. If more than two spots were detected, we calculated the distances between all pairs of spots and considered the largest (maximum metric). The resulting trajectories of distances between detected spots exhibit an oscillatory behavior between the state in which only one spot is visible (compacted state) and the state in which spots are at some distance between each other’s (decompacted state). We define an oscillation as the consecutive time points when more than one spot is detected. From the *mir430* loci trajectories we measured several parameters.

<u>Speed</u>: maximum distance value in oscillation divided by the time it takes to detect only one spot again. For each oscillation in each trajectory, we measured one speed value. We then pooled all speed values and plotted a histogram for all trajectories in presence and absence of Nanog, respectively.

<u>Distances in decompacted state</u>: For each trajectory we measured all distance values for each oscillation using the maximum distance metric as described above. We then pooled all distance values and calculated a histogram for all trajectories in presence and absence of Nanog, respectively.

<u>Time in compacted state (stickiness)</u>: For each trajectory, we measured the duration of all time intervals between two successive oscillations. We then pooled all time values and calculated a histogram for all trajectories in presence and absence of Nanog, respectively.

### Data Normalization

#### Normalization of Nanog intensity

To compare the volume and total intensity of Nanog clusters, the smallest/least intense clusters per nucleus were set to have a value of 0 and the largest/most intense clusters a value of 1. If more than one Nanog cluster colocalized with the same *mir430* transcription body, the volume and total intensity of these Nanog clusters was summed up to obtain only one value per time point and per transcription body. For the rank-based analysis, all Nanog clusters from the same nucleus were ranked based on their total intensity or volume. Percentages of the rank for all Nanog clusters colocalizing with Ser5P transcription bodies (all nuclei) were then plotted in the same density plot or plot for individual nuclei.

To study the evolution of Nanog intensity in the Nanog cluster associated with transcription, we normalized each value to the maximum value of the track. To avoid the bias of very bright Nanog clusters during mitosis, we considered only the values from -150 to +150 seconds around transcription initiation. If one or more spot/shape was detected at one time point, their volume or total intensity were summed up and averaged before normalization.

### Plotting and graph construction

To make graphs, statistics were imported into R-Studio (Integrated Development for R. Rstudio, PBC, Boston, http://www.rstudio.com/). Data were pre-processed and plotted using packages like “ggplot2”, “tidyverse” and “dplyr”.

### Sample size

A minimum of 3 biological replicates (N) was acquired for each experiment. Each biological replicate was obtained from a different and independent batch of embryos. The number of biological replicates (N), embryos, nuclei, and clusters/tracks (n) for each figure panel are given below.

| **Figure** | **Biological replicates (N)** | **Number of embryos** | **Number of nuclei** | **Number of clusters/tracks (n)** |
| --- | --- | --- | --- | --- |
| 1b | 3 | 5 | 25 | 41/3781 |
| 1f | 5/3 | 7/4 | 28/26 | 51 (WT)/48 (inh) |
| 2a | 5 | 7 | 24 | 32 |
| 2b | 5 | 7 | 24 | 32 |
| 2c | 5/4 | 5/6 | 10/13 | 10 (same)/ 13 (different) |
| 2d | 5 | 7 | 28 | 51 |
| 2e | 4/5/5 | 6/7/7 | 16/12/12 | 18 non-merging/13<br>merging (sum) / 26<br>merging (ind) |
| 3d | 3 | 4 | 12 | 24 (WT) |
| 3f | 3 | 4 | 14 | 28 (inh) |
| 4c | 4/4 | 7/7 | 25/38 | 32 (non-merging)/57<br>(merging) |
| 5a, b,c,d,f | 4 | 7 | 46 | 89 |
| 6a, c, d, e | 3/4 | 17/13 | 25/46 | 80 (nanog) / (81<br>no nanog) |
| S2a, b | 3 | 5 | 25 | 41/3781 |
| S3a | 6 | 9 | 36 | / |
| S3b | 4/5/3/3 | 6/7/4/4 | 16/23/13/25 | 18 (WT, non-merging)/ 33<br>(WT, merging)/16<br>(Inh, non-merging)/32<br>(Inh, merging) |
| S5a, b, c, d | 4/4 | 7/7 | 25/38 | 32 (non-merging)<br>/57 (merging) |
| S7b, c, d | 3/3 | 5/5 | 28/22 | 44 (nanog) / (41<br>no nanog) |

### Statistics

P-values for testing the difference in volume and total intensity of *mir430* and other Nanog clusters (Fig. 1b), as well as the difference in time of activation between merging and non-merging clusters (Fig. 2c) were calculated using the paired non-parametric Wilcoxon test. To test the differences between the speed of compaction, duration of compaction and distances in decompacted states between samples with and without Nanog (Fig. 6c, d, e), a one-sided Mann-Whitney test was used.

Because in Figure 6c-e the number of tracks analyzed is small, we performed additional statistical analysis. We randomly split the datasets for each parameter in two smaller populations, both for the data in presence and in absence of Nanog and performed a two-sided Kolmogorow-Smirnow Test (KS-test) to check if they have a similar distribution (data shown below). We report that for all three parameters, this is the case. Moreover, we observed that even when using half of the data, a significant difference in the distribution between the datasets with or without nanog is observed for the time in a compacted state (Figure 6d), and the average maximum distances between densities (Figure 6e), while this is not the case for the speed of compaction (Figure 6c).

|  | **Comparison half-distribution**<br>**With Nanog** | **Comparison half-distribution**<br>**Without Nanog** | **Half with**<br>**Nanog/Half**<br>**without Nanog** |
| --- | --- | --- | --- |
| Figure 6c | p= 0.6872 | p= 0.8617 | p= 0.3076 |
| Figure 6d | p= 0.9017 | p= 0.5861 | p=0.0006171 |
| Figure 6e | p= 0.7046 | p=0.7092 | p= 3.337e-12 |

**Extended Data Figure 1.**
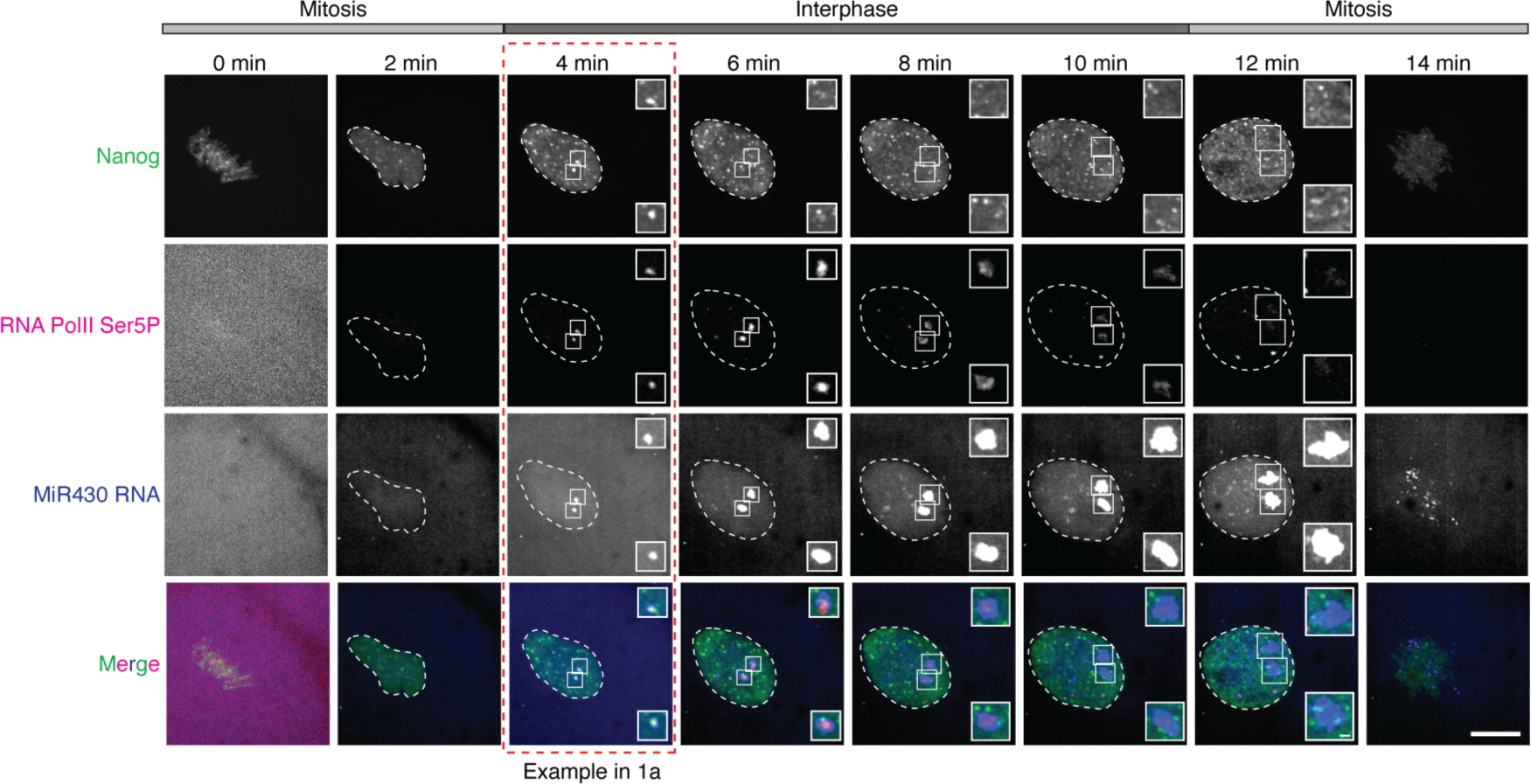
Images of the complete cell cycle for snapshots shown in Fig.1a. Visualization of Nanog (mNeonGreen; green), initiating RNA Polymerase II (RNA Pol II Ser5P Fab (Cy5); magenta), and *mir430* transcription (MoVIE lissamine; blue) during 1k-cell stage. Insets are zooms of the two Nanog clusters colocalizing with RNA Pol II Ser5P and MiR430 transcripts. Shown are all timepoints recorded for an individual nucleus, the time point shown in Figure 1a is boxed in red. Scale bars are 10 and 1 μm (insets). All images represent maximum intensity projections in the z direction.

**Extended Data Figure 2.**
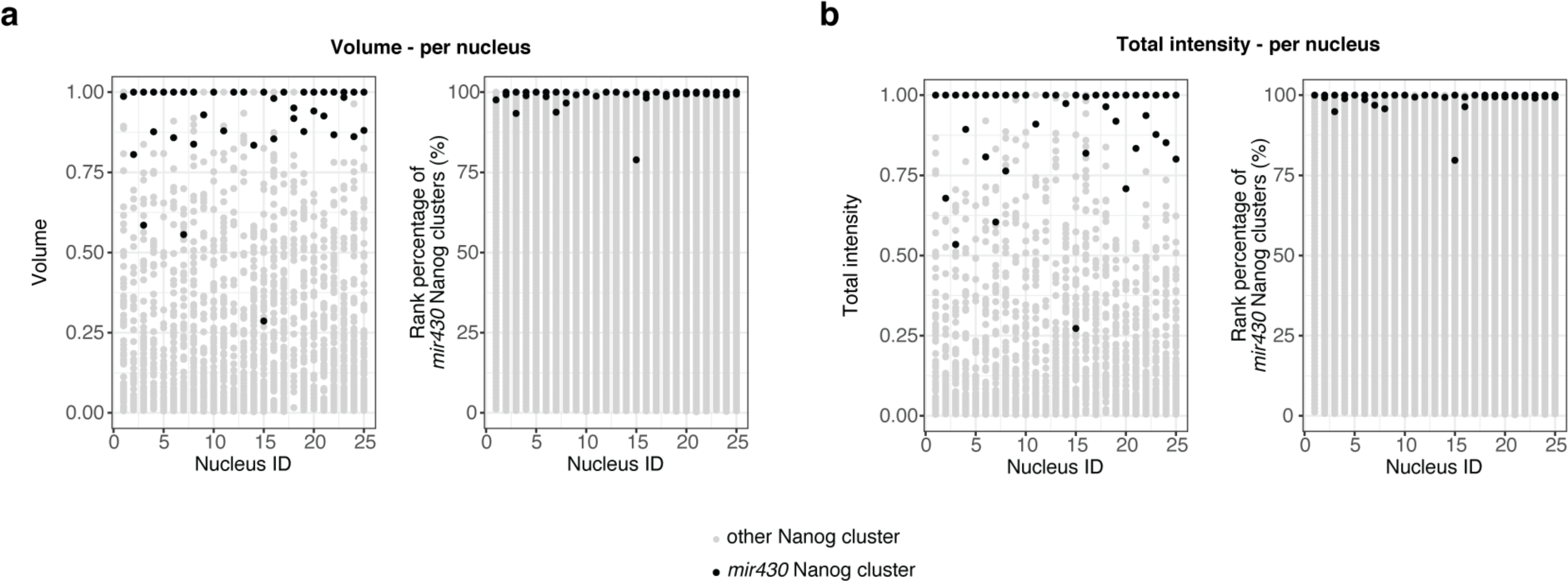
Nanog clusters colocalizing with MiR430 RNA are the largest and brightest in individual nuclei. **a.** Shown are the volume (left), and the rank percentage (right) for Nanog clusters that colocalize with *mir430* transcription (black dots, N=3, n=41) or not (grey dots, N=3, n=3781) in individual nuclei. **b**. Same as in a, but for the total intensity. We note that because the analysis was done at the earliest time-point at which transcription could be detected in a nucleus, and this was sometimes just at one *mir430* allele, there are some nuclei in which just one Nanog cluster was analyzed. For a and b, if two or more Nanog clusters were detected colocalizing with the same RNA Pol II Ser5P transcription body, their volume and total intensity were summed up (see Methods). For a and b, values are normalized for the lowest and the highest values in each nucleus. In this Figure, N is the number of biological replicates, and n is the number of Nanog clusters.

**Extended Data Figure 3.**
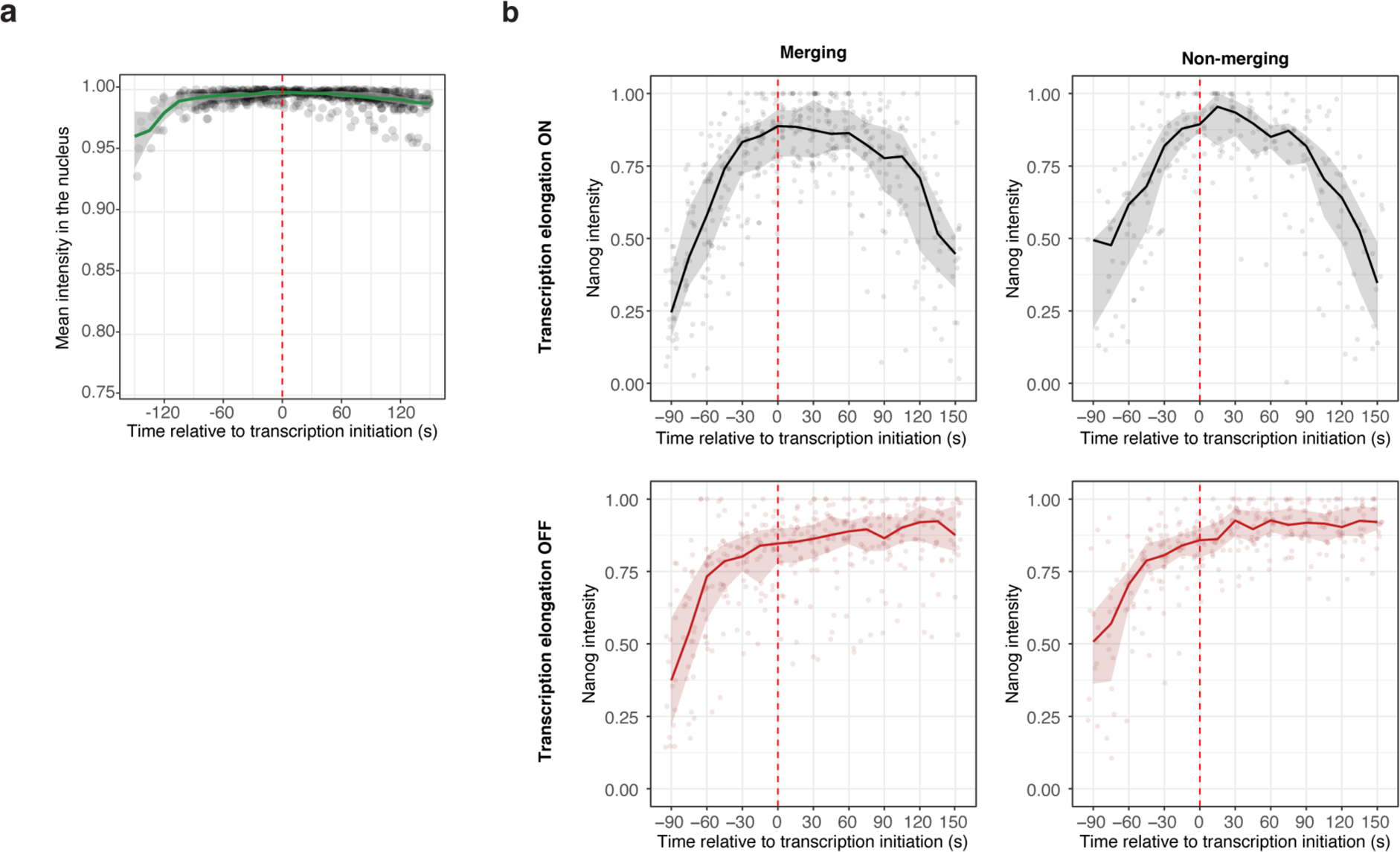
Both merging and non-merging Nanog clusters reach a maximum amount of Nanog independently of transcription elongation. **a.** Mean Nanog nuclear intensity inside the nucleus (without clusters) relative to the time of transcription initiation. Values are normalized to the maximum value for the same nucleus. Green line indicates the median of the distribution and grey ribbons respectively 25% and 75% of the distribution for the lower and upper limits at each time point. **b.** Related to Fig. 1f. Total intensity of Nanog clusters associated with the *mir430* DNA locus relative to the time of transcription initiation, with (black) and without (red) transcription elongation, split between merging and non-merging clusters (non-merging, no inhibition, N=4, n=16; non-merging, transcription inhibition, N=5, n=23; merging, no inhibition, N=3, n=19; merging, transcription inhibition, N=3, n=19). If the Nanog cluster associated with transcription was the result of a merging event, we summed up the total intensity of all clusters per time point. The bold line represents the median and ribbon the 25^th^ and 75^th^ percentile of the distribution. Values are normalized to the maximum value for each track. The red dash line indicates transcription initiation. In this Figure, N is the number of biological replicates, and n is the number of Nanog clusters.

**Extended Data Figure 4.**
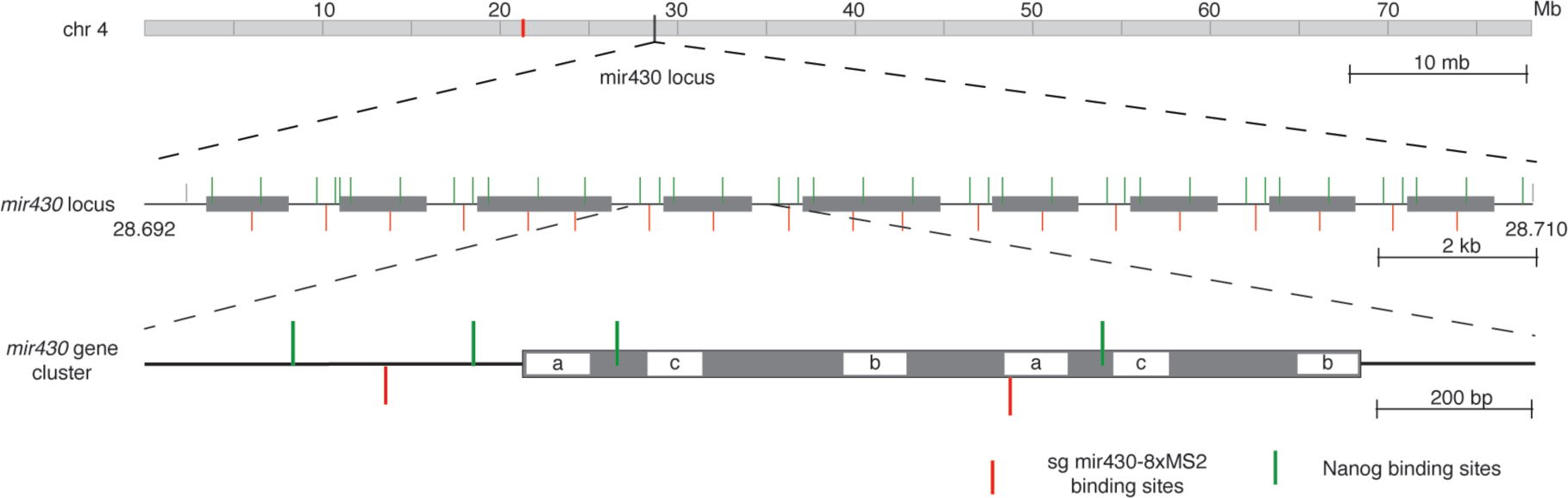
Structure of the *mir430* locus. Shown is the structure of the *mir430* locus on the long arm of chromosome IV, as described in the GRCz11 zebrafish genome assembly (Howe et al., 2013). The three isoforms of *mir430* are indicated by the labels ‘a’,’b’ and ‘c’. Red and green bars indicate binding sites for the *mir430* sgRNA and predicted Nanog motifs, respectively.

**Extended Data Figure 5.**
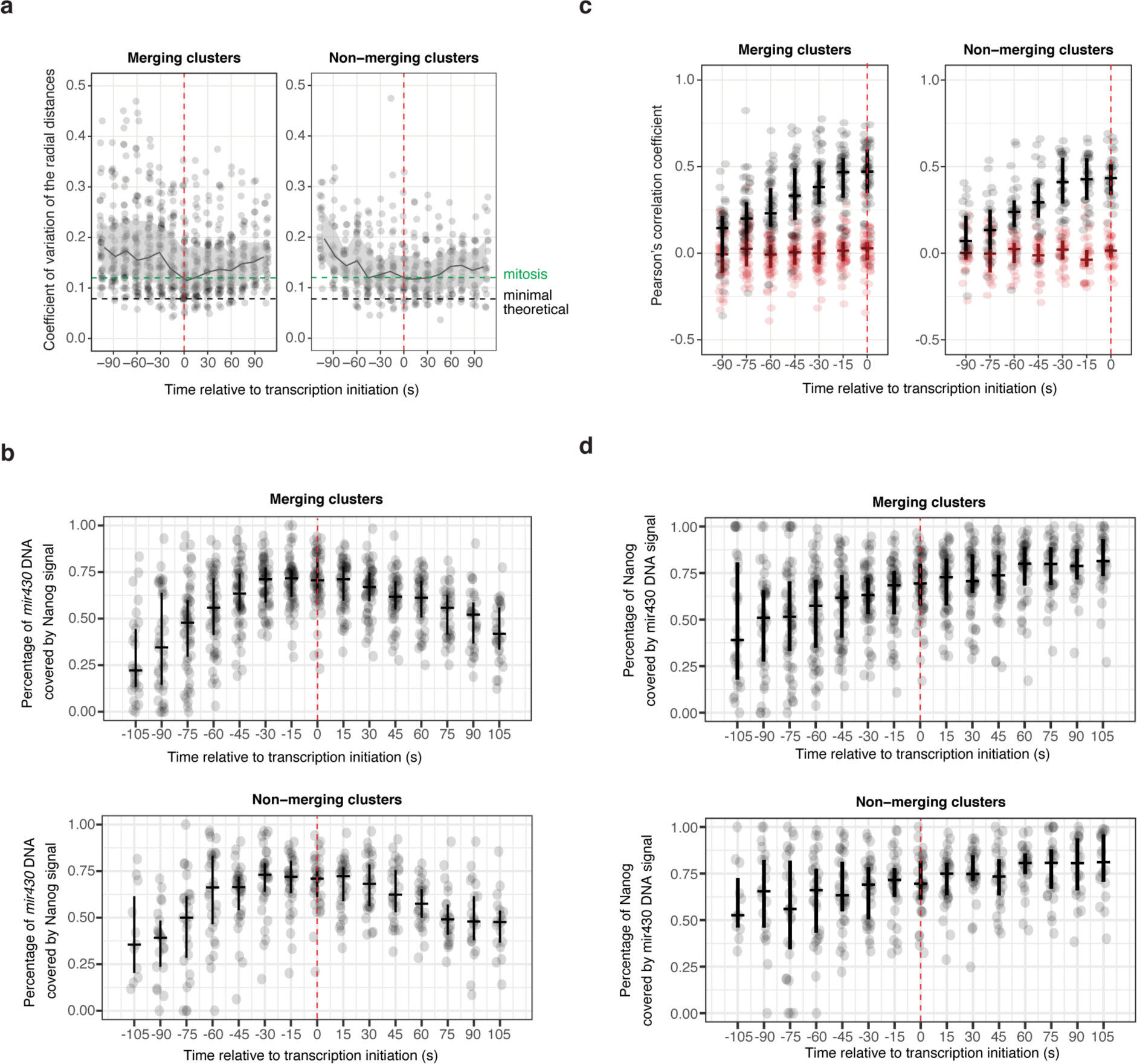
Merging and non-merging cases display similar behavior. **a.** Related to Fig. 5a, here separated for merging and non-merging clusters. The CoV of the radial distances for all tracks is plotted as a function of time, centered on transcription initiation. The grey line indicates the median of the distribution and the associated ribbon the 25th and 75th percentile of the distribution. The green dashed line shows the average value of the coefficient of variation of the radial distance of the *mir430* DNA mask measured during mitosis (see Methods). The black dashed line shows the theoretical minimal value for a mask considered as a perfect sphere. **b.** Related to Fig. 5b, here separated for merging and non-merging clusters. Percentage of *mir430* DNA signal that is covered by Nanog signal as a function of time. The vertical lines represent respectively the 25 and 75% of the distribution, while the horizontal line represents the median. **c.** Related to Fig. 5c, here separated for merging and non-merging clusters. Boxplots showing the Pearson’s correlation score between *mir430* DNA mask and the associated Nanog signal (black) or scrambled Nanog signal (red) for all tracks, relative to the start of transcription. **d.** Related to Fig. 5d, here separated for merging and non-merging clusters. Percentage of *mir430*-associated Nanog signal that is covered by *mir430* DNA signal as a function of time. For all panels, the red dash line indicates transcription initiation. In this Figure, with N=biological replicates and n=*mir430* alleles, N=4 and n=32 for non-merging clusters, and N=4 and n=57 for merging clusters.

**Extended Data Figure 6.**
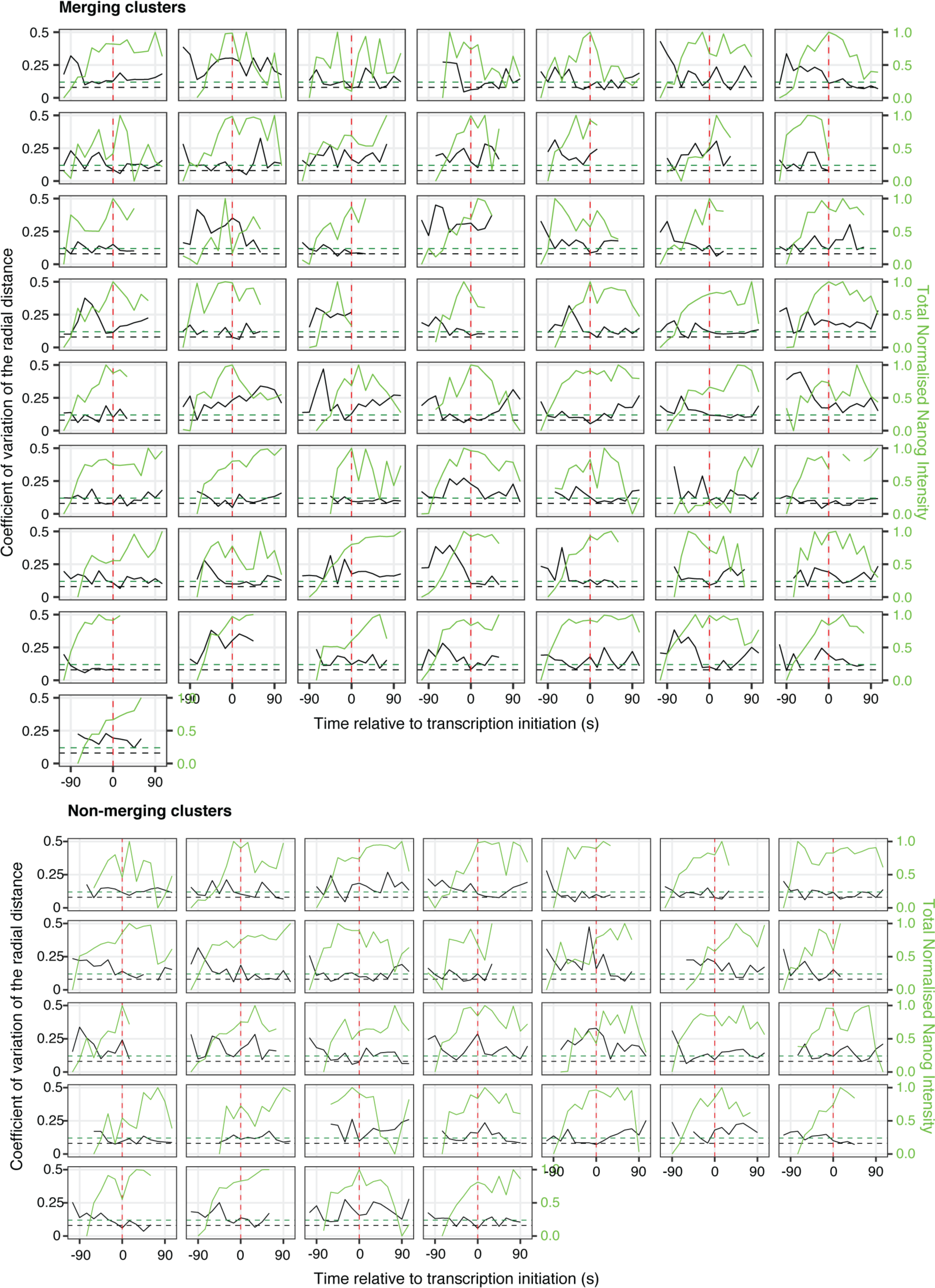
Individual plots for data shown in Fig. 5. Coefficient of variation of radial distances of single *mir430* DNA alleles, as well as total intensity of associated Nanog signal as a function of time, relative to transcription initiation (indicated with a red dashed line). Green dashed line represents the average value of the coefficient of variation of radial distances of the *mir430* DNA mask during mitosis. The black dashed line shows the minimal value for a mask considered as a perfect sphere.

**Extended Data Figure 7.**
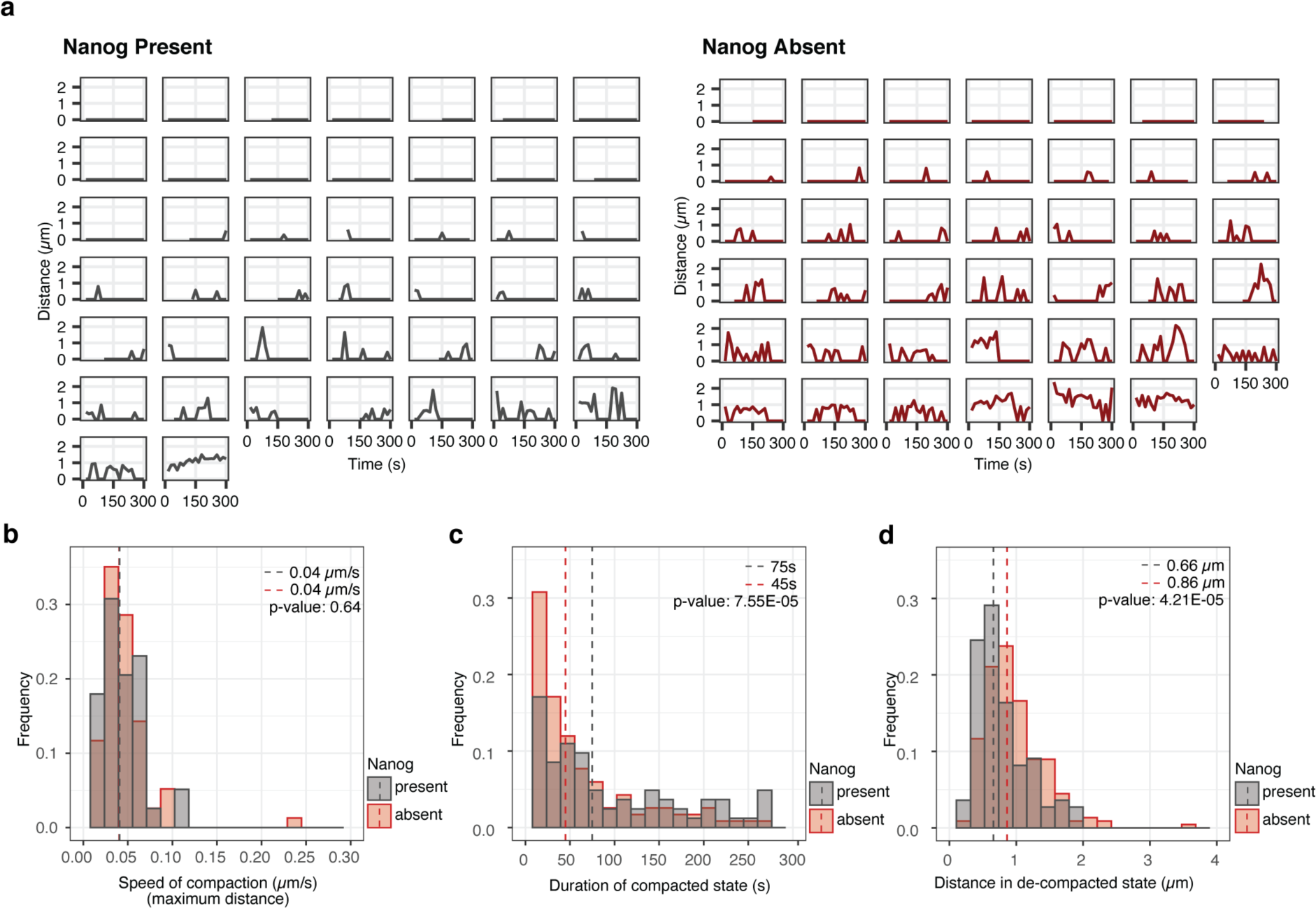
1k-cell stage data from Fig. 6. **a**. Graphs of the distances between detected densities on the *mir430* DNA channel as a function of time for individual *mir430* DNA alleles in the presence (black) or absence (red) at 1k-cell stage only. **b-d.** Histograms showing the speed of compaction (from the maximum distance) (c), the duration of compacted states (or stickiness) (d), and the distances in decompacted states (e) in the presence (black, N=3 and n=44) and absence (red, N=3 and n=41) of Nanog for the 1k-cell stage. Dashed lines indicate the medians of the distributions. P-values are calculated with one-sided Mann-Whitney test. In this Figure, N is the number of biological replicates, and n is the number of *mir430* alleles.

**Supplementary Video 1. Movie of Nanog clusters merging followed by transcription initiation. Related to** Figure 1c. Spinning-disk confocal microscope time-lapse of Nanog clusters (mNeonGreen; left) and RNA Pol II Ser5P (RNA Pol II Ser5P Fab (Cy5); right) at 1k-cell stage. Yellow and cyan arrowheads on the left point to merging and non-merging clusters, respectively. White arrowheads on the right represent the transcription bodies. Time is relative to transcription initiation. All images are snapshots from the 3D rendering of the Imaris software.

**Supplementary Video 2. Complete movie of merging Nanog clusters with associated *mir430* DNA and MiR430 RNA. Related to** Figure 4b **(i).** Spinning-disk confocal microscope time-lapse of *mir430* DNA (tdMCP-mNG; green), MiR430 RNA (MoVIE-lissamine; blue) and Nanog (HaloTag (JFX650); magenta) in *nanog* -/- embryos at 1k-cell stage. Yellow arrowheads on the left point to the two merging Nanog clusters and on the right to the associated transcription body. Time is relative to transcription initiation. All images are snapshots from the 3D rendering of the Imaris software.

**Supplementary Video 3. Complete movie of Nanog clusters rapidly splitting and merging with associated *mir430* DNA and MiR430 RNA. Related to** Figure 4b **(ii).** Spinning-disk confocal microscope time-lapse of *mir430* DNA (tdMCP-mNG; green), MiR430 RNA (MoVIE-lissamine; blue) and Nanog (HaloTag (JFX650); magenta) in *nanog* -/- embryos at 1k-cell stage. Yellow arrowheads on the left point to merging Nanog clusters and on the right to the associated transcription body. Time is relative to transcription initiation. All images are snapshots from the 3D rendering of the Imaris software.

**Supplementary Video 4. Complete movie of a unique Nanog cluster with associated *mir430* DNA and MiR430 RNA. Related to** Figure 4b **(iii).** Spinning-disk confocal microscope time-lapse of *mir430* DNA (tdMCP-mNG; green), MiR430 RNA (MoVIE-lissamine; blue) and Nanog (HaloTag (JFX650); magenta) in *nanog* -/- embryos at 1k-cell stage. Yellow arrowheads on the left point to the non-merging Nanog cluster and on the right to the associated transcription body. Time is relative to transcription initiation. All images are snapshots from the 3D rendering of the Imaris software.

## References

1. Cho, W.-K. et al. Mediator and RNA polymerase II clusters associate in transcription-dependent condensates. Science 361, 412–415 (2018).

2. Kuznetsova, K. et al. Nanog organizes transcription bodies. Curr. Biol. 33, 164–173.e5 (2023).

3. Li, J. et al. Single-gene imaging links genome topology, promoter–enhancer communication and transcription control. Nat. Struct. Mol. Biol. 27, 1032–1040 (2020).

4. Mir, M. et al. Dynamic multifactor hubs interact transiently with sites of active transcription in Drosophila embryos. eLife 7, (2018).

5. Nair, S. J. et al. Phase separation of ligand-activated enhancers licenses cooperative chromosomal enhancer assembly. Nat. Struct. Mol. Biol. 2019 263 26, 193–203 (2019).

6. Sabari, B. R. et al. Coactivator condensation at super-enhancers links phase separation and gene control. Science 361, (2018).

7. Tsai, A. et al. Nuclear microenvironments modulate transcription from low-affinity enhancers. eLife 6, (2017).

8. Tsai, A., Alves, M. R. P. & Crocker, J. Multi-enhancer transcriptional hubs confer phenotypic robustness. eLife 8, (2019).

9. Hyman, A. A., Weber, C. A. & Jülicher, F. Liquid-Liquid Phase Separation in Biology. Annu. Rev. Cell Biol 30, 39–58 (2014).

10. Kent, S. et al. Phase-Separated Transcriptional Condensates Accelerate Target-Search Process Revealed by Live-Cell Single-Molecule Imaging. Cell Rep. 33, 108248 (2020).

11. Ahn, J. H. et al. Phase separation drives aberrant chromatin looping and cancer development. Nature 595, 591–595 (2021).

12. Chowdhary, S., Kainth, A. S., Paracha, S., Gross, D. S. & Pincus, D. Inducible transcriptional condensates drive 3D genome reorganization in the heat shock response. Mol. Cell 82, 4386–4399.e7 (2022).

13. Wang, W. et al. A histidine cluster determines YY1-compartmentalized coactivators and chromatin elements in phase-separated enhancer clusters. Nucleic Acids Res. 50, 4917–4937 (2022).

14. Ma, L. et al. Co-condensation between transcription factor and coactivator p300 modulates transcriptional bursting kinetics. Mol. Cell 81, 1682–1697.e7 (2021).

15. Wei, M. T. et al. Nucleated transcriptional condensates amplify gene expression. Nat. Cell Biol. 22, 1187–1196 (2020).

16. Chen, L. et al. Hormone-induced enhancer assembly requires an optimal level of hormone receptor multivalent interactions. Mol. Cell 83, 3438–3456.e12 (2023).

17. Kim, Y. J. et al. Light-activated macromolecular phase separation modulates transcription by reconfiguring chromatin interactions. Sci. Adv. 9, (2023).

18. Schneider, N. et al. Liquid-liquid phase separation of light-inducible transcription factors increases transcription activation in mammalian cells and mice. Sci. Adv. 7, (2021).

19. Wu, J. et al. Modulating gene regulation function by chemically controlled transcription factor clustering. Nat. Commun. 13, (2022).

20. Du, M. et al. Direct observation of a condensate effect on super-enhancer controlled gene bursting. Cell 187, 331–344.e17 (2024).

21. Kawasaki, K. & Fukaya, T. Functional coordination between transcription factor clustering and gene activity. Mol. Cell 83, 1605–1622.e9 (2023).

22. Song, L. et al. Hotspot mutations in the structured ENL YEATS domain link aberrant transcriptional condensates and cancer. Mol. Cell 82, 4080–4098.e12 (2022).

23. Li, J. et al. Single-Molecule Nanoscopy Elucidates RNA Polymerase II Transcription at Single Genes in Live Cells. Cell 178, 491–506.e28 (2019).

24. Meeussen, J. V. W. et al. Transcription factor clusters enable target search but do not contribute to target gene activation. Nucleic Acids Res. 51, (2023).

25. Xie, J. et al. Targeting androgen receptor phase separation to overcome antiandrogen resistance. Nat. Chem. Biol. 18, 1341–1350 (2022).

26. Zhang, H. et al. Reversible phase separation of HSF1 is required for an acute transcriptional response during heat shock. Nat. Cell Biol. 24, 340–352 (2022).

27. Furlong, E. E. M. & Levine, M. Developmental enhancers and chromosome topology. Science 361, 1341–1345 (2018).

28. Hnisz, D., Shrinivas, K., Young, R. A., Chakraborty, A. K. & Sharp, P. A. Leading Edge Perspective A Phase Separation Model for Transcriptional Control. Cell 169, 13–23 (2017).

29. Shrinivas, K. et al. Enhancer Features that Drive Formation of Transcriptional Condensates In Brief Transcription-associated proteins form condensates localized at specific DNA elements. Mol. Cell 75, 549–561.e7 (2019).

30. Saravanan, B. et al. Ligand dependent gene regulation by transient ERα clustered enhancers. PLoS Genet. 16, e1008516 (2020).

31. Whyte, W. A. et al. Master transcription factors and mediator establish super-enhancers at key cell identity genes. Cell 153, 307–319 (2013).

32. Keenen, M. M. et al. HP1 proteins compact DNA into mechanically and positionally stable phase separated domains. eLife 10, e64563 (2021).

33. Nguyen, T. et al. Chromatin sequesters pioneer transcription factor Sox2 from exerting force on DNA. Nat. Commun. 13, 3988 (2022).

34. Quail, T. et al. Force generation by protein–DNA co-condensation. Nat. Phys. 2021 179 17, 1007–1012 (2021).

35. Heyn, P. et al. The earliest transcribed zygotic genes are short, newly evolved, and different across species. Cell Rep. 6, 285–292 (2014).

36. White, R. J. et al. A high-resolution mRNA expression time course of embryonic development in zebrafish. eLife 6, (2017).

37. Chan, S. H., Tang, Y., Bazzini, A. A., Moreno-Mateos, M. A. & Giraldez, A. J. Brd4 and P300 Confer Transcriptional Competency during Zygotic Genome Activation. Dev. Cell 49, 867–881.e8 (2019).

38. Hadzhiev, Y. et al. A cell cycle-coordinated Polymerase II transcription compartment encompasses gene expression before global genome activation. Nat. Commun. 10, (2019).

39. Hadzhiev, Y. et al. The miR-430 locus with extreme promoter density forms a transcription body during the minor wave of zygotic genome activation. Dev. Cell 58, 155–170.e8 (2023).

40. Hilbert, L. et al. Transcription organizes euchromatin via microphase separation. Nat. Commun. 2021 121 12, 1–12 (2021).

41. Pownall, M. E. et al. Chromatin expansion microscopy reveals nanoscale organization of transcription and chromatin. Science 381, 92 (2023).

42. Ugolini, M. et al. Transcription bodies regulate gene expression by sequestering CDK9. Nat. Cell Biol. 26, 604–612 (2024).

43. Lee, M. T. et al. Nanog, Pou5f1 and SoxB1 activate zygotic gene expression during the maternal-to-zygotic transition. Nature 503, 360–364 (2013).

44. Hayashi-Takanaka, Y. et al. Tracking epigenetic histone modifications in single cells using Fab-based live endogenous modification labeling. Nucleic Acids Res. 39, 6475–6488 (2011).

45. Kimura, H. & Yamagata, K. Visualization of epigenetic modifications in preimplantation embryos. Methods Mol. Biol. 1222, 127–147 (2015).

46. Sato, Y. et al. Histone H3K27 acetylation precedes active transcription during zebrafish zygotic genome activation as revealed by live-cell analysis. Development 146, 19 (2019).

47. Stasevich, T. J. et al. Regulation of RNA polymerase II activation by histone acetylation in single living cells. Nat. 2014 5167530 516, 272–275 (2014).

48. Rudd, M. D. & Luse, D. S. Amanitin greatly reduces the rate of transcription by RNA polymerase II ternary complexes but fails to inhibit some transcript cleavage modes. J. Biol. Chem. 271, 21549–21558 (1996).

49. Dominguez, A. A., Lim, W. A. & Qi, L. S. Beyond editing: repurposing CRISPR–Cas9 for precision genome regulation and interrogation. Nat. Rev. Mol. Cell Biol. 17, 5–15 (2016).

50. Ma, H. et al. CRISPR-Sirius: RNA scaffolds for signal amplification in genome imaging. Nat. Methods 15, 928–931 (2018).

51. Leidescher, S. et al. Spatial organization of transcribed eukaryotic genes. Nat. Cell Biol. 24, 327– 339 (2022).

52. Chong, S. et al. Tuning levels of low-complexity domain interactions to modulate endogenous oncogenic transcription. Mol. Cell 82, 2084–2097.e5 (2022).

53. Mazzocca, M., Fillot, T., Loffreda, A., Gnani, D. & Mazza, D. The needle and the haystack: single molecule tracking to probe the transcription factor search in eukaryotes. Biochem. Soc. Trans. 49, 1121 (2021).

54. Trojanowski, J. et al. Transcription activation is enhanced by multivalent interactions independent of phase separation. Mol. Cell 82, 1878–1893.e10 (2022).

55. Xu, C. et al. Nanog-like Regulates Endoderm Formation through the Mxtx2-Nodal Pathway. Dev. Cell 22, 625 (2012).

56. Boija, A. A. et al. Transcription Factors Activate Genes through the Phase-Separation Capacity of Their Activation Domains In Brief Activation domains from a diverse array of mammalian and yeast transcription factors form phase-separated condensates with Mediator to activate gene expression. Cell 175, 1842–1855.e16 (2018).

57. Chong, S. et al. Imaging dynamic and selective low-complexity domain interactions that control gene transcription. Science 361, (2018).

58. Shi, B. et al. UTX condensation underlies its tumour-suppressive activity. Nature 597, 726–731 (2021).

59. Heurtier, V. et al. The molecular logic of Nanog-induced self-renewal in mouse embryonic stem cells. Nat. Commun. 10, 1109 (2019).

60. Silva, J. et al. Nanog Is the Gateway to the Pluripotent Ground State. Cell 138, 722–737 (2009).

61. Theunissen, T. W. et al. Nanog Overcomes Reprogramming Barriers and Induces Pluripotency in Minimal Conditions. Curr. Biol. 21, 65–71 (2011).

62. Allègre, N. et al. NANOG initiates epiblast fate through the coordination of pluripotency genes expression. Nat. Commun. 13, 3550 (2022).

63. Piazzolla, D. et al. Lineage-restricted function of the pluripotency factor NANOG in stratified epithelia. Nat. Commun. 5, 4226 (2014).

64. Wang, Z., Oron, E., Nelson, B., Razis, S. & Ivanova, N. Distinct lineage specification roles for NANOG, OCT4, and SOX2 in human embryonic stem cells. Cell Stem Cell 10, 440–454 (2012).

65. Miao, L. et al. The landscape of pioneer factor activity reveals the mechanisms of chromatin reprogramming and genome activation. Mol. Cell 82, 986–1002.e9 (2022).

66. Pálfy, M., Schulze, G., Valen, E. & Vastenhouw, N. L. Chromatin accessibility established by Pou5f3, Sox19b and Nanog primes genes for activity during zebrafish genome activation. PLoS Genet. 16, e1008546 (2020).

67. Veil, M., Yampolsky, L. Y., Grüning, B. & Onichtchouk, D. Pou5f3, SoxB1, and Nanog remodel chromatin on high nucleosome affinity regions at zygotic genome activation. Genome Res. 29, 383–395 (2019).

68. Veil, M. et al. Maternal Nanog is required for zebrafish embryo architecture and for cell viability during gastrulation. (2018) doi:10.1242/dev.155366.

69. The Zebrafish Book: A Guide for the Laboratory Use of Zebrafish (Danio Rerio) - Monte Westerfield - Google Books.

70. Waldo, G. S., Standish, B. M., Berendzen, J. & Terwilliger, T. C. Rapid protein-folding assay using green fluorescent protein. Nat. Biotechnol. 17, 691–695 (1999).

71. Hayashi-Takanaka, Y. et al. Tracking epigenetic histone modifications in single cells using Fab-based live endogenous modification labeling. Nucleic Acids Res. 39, 6475–6488 (2011).

72. Schindelin, J. et al. Fiji: an open-source platform for biological-image analysis. Nat. Methods 2012 97 9, 676–682 (2012).

73. Ershov, D. et al. TrackMate 7: integrating state-of-the-art segmentation algorithms into tracking pipelines. Nat. Methods 19, 829–832 (2022).

